# gEL DNA, a cloning- and PCR-free method for CRISPR-based multiplexed genome editing

**DOI:** 10.1101/2020.05.22.110494

**Authors:** Paola Randazzo, Jean-Marc Daran, Pascale Daran-Lapujade

## Abstract

Even for the genetically accessible yeast *Saccharomyces cerevisiae*, the CRISPR/Cas DNA editing technology has strongly accelerated and facilitated strain construction. Several methods have been validated for fast and highly efficient single editing events and diverse approaches for multiplex genome editing have been described in literature by means of Cas9 or Cas12a endonucleases and their associated gRNAs. The gRNAs used to guide the Cas endonuclease to the editing site are typically expressed from plasmids using native PolII or PolIII RNA polymerases. These gRNA-expression plasmids require laborious, time-consuming cloning steps, which hampers their implementation for academic and applied purposes. In this study, we explore the potential of expressing gRNA from linear DNA fragments using the T7 RNA polymerase (T7RNAP) for single and multiplex genome editing in *S. cerevisiae*. Using Cas12a, this work demonstrates that transforming short, linear DNA fragments encoding gRNAs in yeast strains expressing T7RNAP promotes highly efficient single DNA editing. These DNA fragments can be custom-ordered, which makes this approach highly suitable for high-throughput strain construction. This work expands the CRISPR-toolbox for large-scale strain construction programs in *S. cerevisiae* and promises to be relevant for other, less genetically accessible yeast species.

## Introduction

The bacterial-derivative CRISPR-Cas technology is nowadays the most commonly used tool for microbial genome engineering. For the eukaryotic model and industrial workhorse *Saccharomyces cerevisiae*, several CRISPR-based methodologies have been developed aiming at fast and efficient single editing event (1–4). Two Class II bacterial endonucleases, Cas9 and Cas12a (also known as Cpf1) have been functionally characterized for DNA editing ranging from point mutation to heterologous pathway integration (4–6). While diverse Cas9- and Cas12a-mediated approaches for multiplex genome editing have been described in literature (reviewed in (7)), multiplex genome editing still requires substantial efforts for the CRISPR tools to be built. The RNA molecules designed to guide the endonuclease towards the editing site (gRNAs) are typically cloned in and expressed from plasmids. In most published works so far, multiplex editing relies on the parallel transformation of multiple plasmids carrying a single or two gRNAs. However, this approach is limited by the number of available marker-based plasmid backbones (4,8–12). More recently, several successful examples have shown that several gRNAs can be expressed from a single gRNA-array, using different tricks to release the mature gRNAs (3,5,6,13–17). However, complexity of these gRNA expression cassettes and their tailored sequence design may be difficult to synthesize and requires laborious and time-consuming cloning steps, therefore hindering the workflow for strain construction. To date, few attempts have been developed to circumvent gRNA cloning for genome editing of microbes in general and of *S. cerevisiae* in particular.

The most straightforward, cloning-free strategy would rely on the delivery of the gRNA in the form of a short, linear DNA fragment. Such short DNA fragments could easily be synthetized as oligonucleotides and delivered as mixture in any desired gRNA combination for multiplex targeting of DNA sites. Such a cost-effective and versatile approach would be highly suited for high-throughput, multiplex genome engineering of strains. Transient expression of linear DNA carrying gRNA expression cassettes has been previously shown to enable Cas9-mediated DNA-editing (9,18). However, these approaches systematically require a first *in vitro* step for the construction of vectors from which the linear DNA is produced by PCR amplification (Fig. 1). In eukaryotes, gRNAs are transcribed either by RNA polymerase III (RNApolIII) or by RNA Polymerase II (RNAPolII) promoter, this latter being flanked by self-processing ribozymes or tRNAs that prevent unwanted processing of the gRNAs (16,17). A recent report has shown that functional gRNAs can also be transcribed in different yeasts by the RNA polymerase from bacteriophage T7 (T7RNAP) localized in the nucleus (19). Delivered as plasmid DNA, the T7RNAP-transcribed gRNAs have been used to guide Cas9 for genome editing and dCas9 for transcriptional regulation.

**Figure 1:**
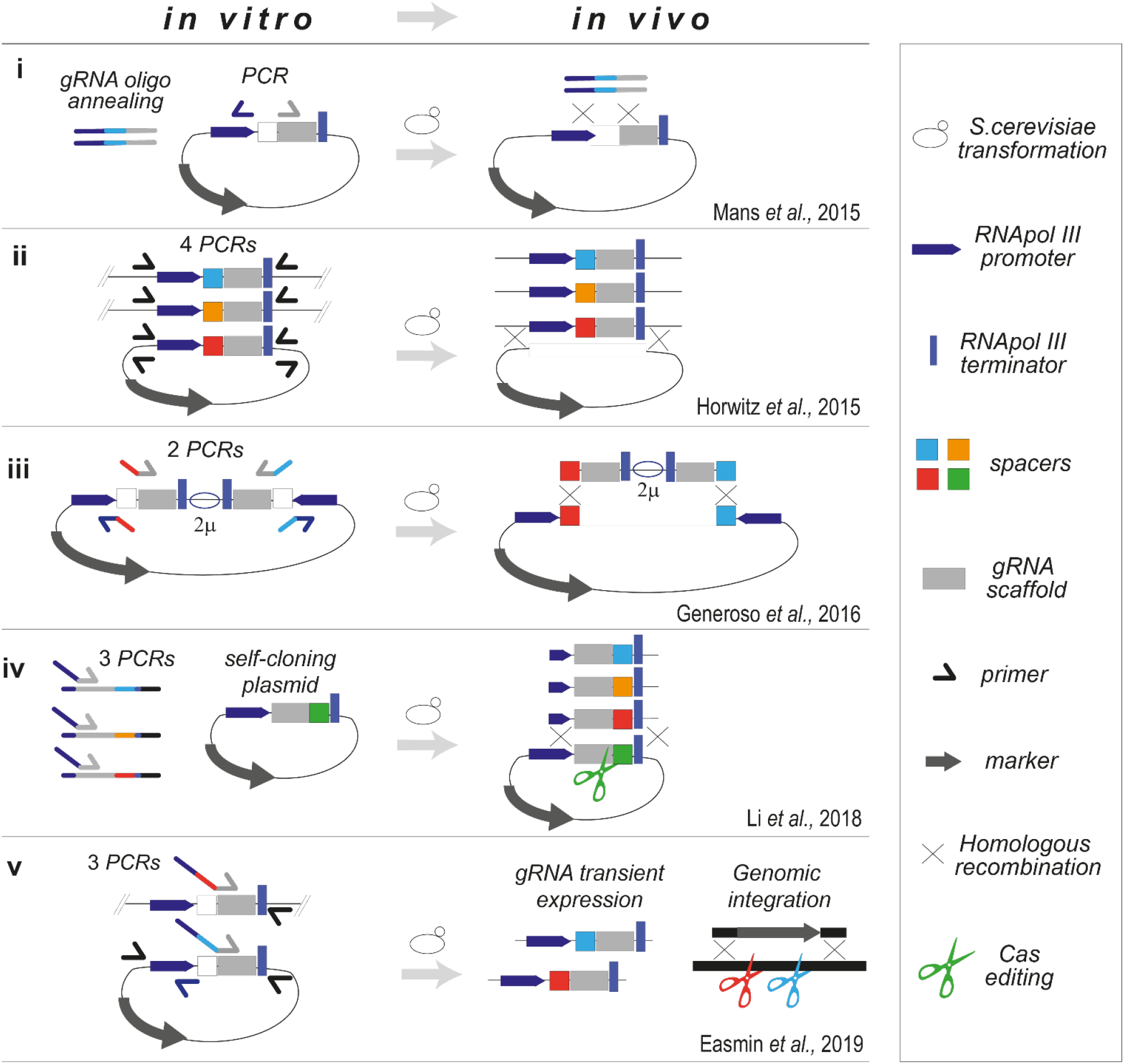
cloning-free approaches for CRISPR/Cas-aided DNA editing. Overview of methodologies based on delivery of linear DNA templates for gRNAs expression in *S. cerevisiae*. *In vitro* sample preparations and *in vivo* events upon transformation are described. Number of PCR reactions are quoted. Features are depicted in the legend at the right handside of the figure.

The present work introduces the <u>gEL DNA</u> method, a novel, utterly cloning and PCR-free genome editing tool, based on the <u>g</u>RNA <u>E</u>xpression from short, <u>L</u>inear double-stranded <u>DNA</u> oligos by the T7RNAP (Fig. 2). Comparing Cas9 and Cas12a, this study demonstrates that Cas12a enables efficient single and multiplexed DNA editing from custom-ordered oligonucleotides of 87 nt in *S. cerevisiae*. Next to gRNA *in silico* design, the only steps required for genome editing are transformation and screening. Highly suited for high-throughput strain construction, the gEL DNA method does not require prior knowledge on the transcription machinery of the host microbe (e.g. RNA processing and promoters) and thereby promises to facilitate DNA editing in less genetically accessible microbes.

**Figure 2:**
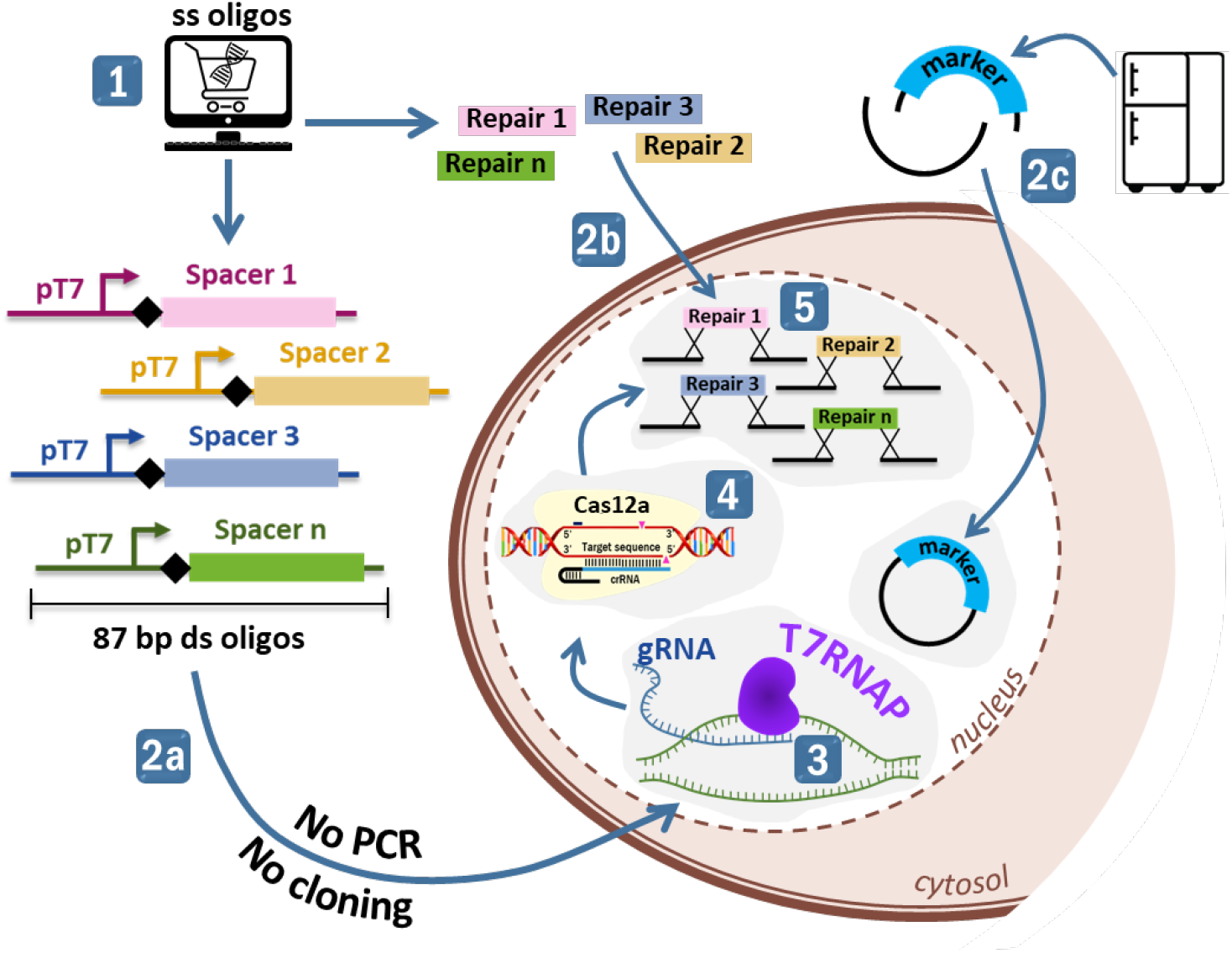
Schematic overview of the gEL DNA approach. 1, *in silico* design and ordering of gDNA cassettes (87 bp) and repair DNA (120 bp) as oligos. 2, tranformation with the double-stranded (ds) gDNA expression cassettes (2a), the ds repair DNA fragments and an empty, split plasmid carrying a marker of choice. 3, expression of the gRNA by the T7RNAP. 4, targetted DNA editing by Cas12a (Cas12a). 5, repair of the ds DNA break via homologous recombination using the repair DNA fragments.

## Materials and Methods

### Strains and cultivation conditions

All *S. cerevisiae* strains used in this study (Table 1) were derived from the CEN.PKbackground strain (20). Yeast cells were grown at 30 °C in shake flasks on rotary shaker (200 rpm) or on agar plates (20 g l^−1^). Complex medium contained 10 g l^−1^ of yeast extract, 20 g l^−1^ of peptone and 20 g l^−1^ of glucose (YPD). YPD was supplemented with nourseothricin (100 mg l^−1^), geneticin (G418) (200 mg l^−1^) or hygromycin B (200 mg l^−1^) to select transformants. Minimal synthetic media were prepared as previously described (21). SMD medium contained 5 g l^−1^ of (NH_4_)_2_SO_4_, 3 g l^−1^ of KH_2_PO_4_, 0.5 g l^−1^ of MgSO_4_·7H_2_O, 1 ml l^−1^ of a trace element solution, supplemented with 20 g l^−1^ of glucose and 1 ml l^−^ of a vitamin solution. SMD-urea included 6.6 g l^−1^ K_2_SO_4_, 3.0 g·l^−1^ KH_2_PO_4_, 0.5 g l^−1^ MgSO_4_·7H_2_O, 1 mL l^−1^ trace elements solution, supplemented with 20 g l^−1^ of glucose, 1 ml l^−1^ of a vitamin solution and 2.3 g l^−1^ CH_4_N_2_O (22). For selection of transformants carrying the *amdS* marker cassette, ammonium sulfate in SMD was substituted with 10 mM acetamide and 6.6 g L^−1^ K_2_SO_4_ (SM-Ac) (23). Plasmids were propagated in *Escherichia coli* XL1-Blue cells (Agilent Technologies, Santa Clara, CA), after growth in Lysogeny broth (LB; 10 g l^−1^ tryptone, 5 g l^−1^ yeast extract, 10 g l^−1^ NaCl) liquid culture (180 rpm) or solid medium (20 g l^−1^ agar) supplemented with chloramphenicol (25 mg l^−1^), spectinomycin (100 mg l^−1^) or ampicillin (100 mg l^−1^) at 37 °C. When required, plasmids from yeasts isolates were removed accordingly to described procedures (4). All *S. cerevisiae* and *E. coli* stocks were prepared by aseptically adding 30% v/v of glycerol to exponentially growing cultures. Aliquoted cell stocks were stored at −80 °C.

**Table 1 –.**
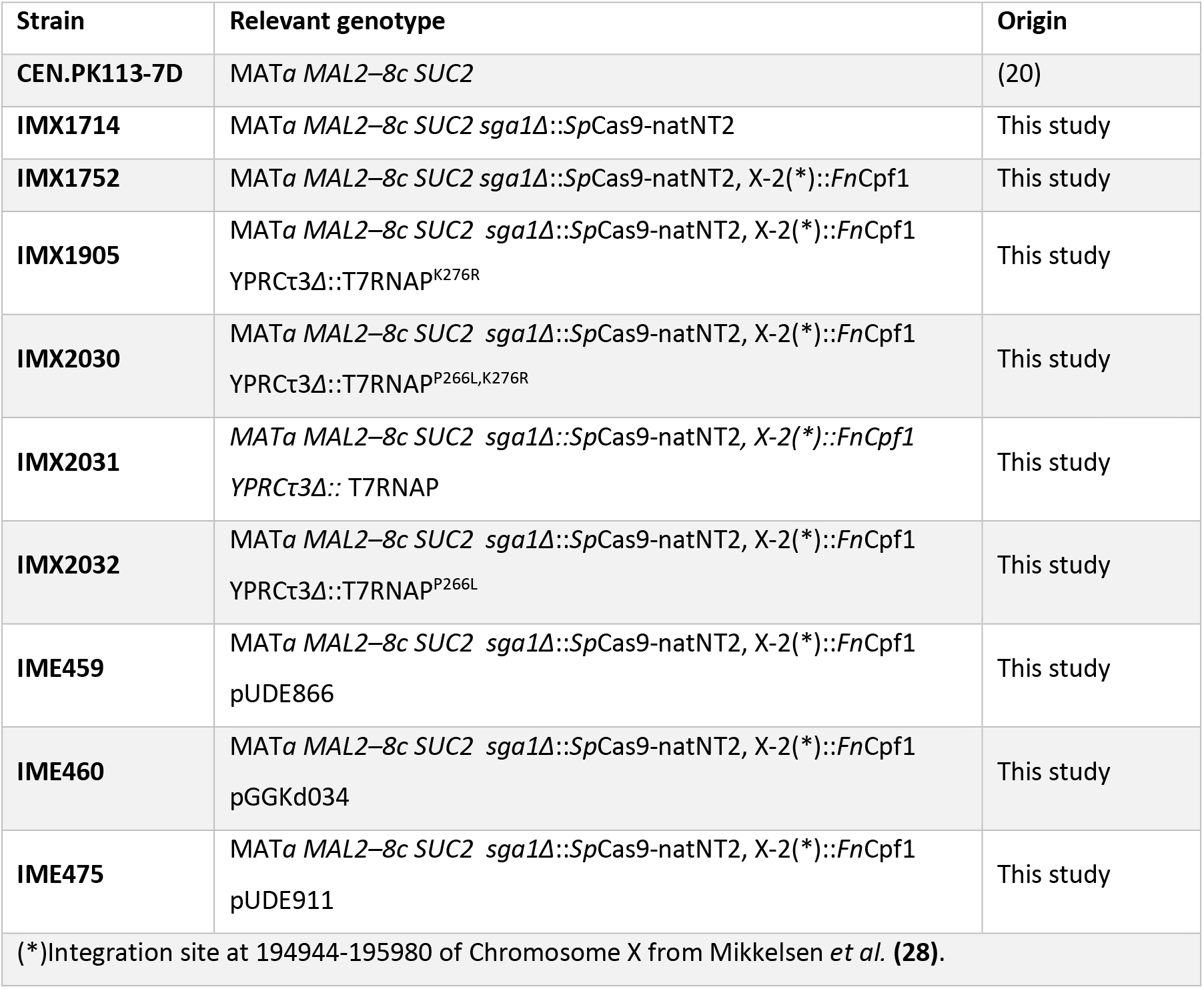
*Saccharomyces cerevisiae* strains used in this study

### Molecular biology techniques

Yeast genomic DNA used for cloning purposes was isolated using the YeaStar genomic DNA kit (Zymo Research, Irvine, CA) according to manufacturer’s instructions. Diagnostic PCR was performed using DreamTaq DNA polymerase (Thermo Fisher Scientific, Walthman, MA). For cloning and sequencing purposes, PCR products were obtained using Phusion^®^ High-Fidelity DNA Polymerase (Thermo Fisher Scientific). Primers were ordered as PAGE or desalted purified oligonucleotides (Table S1) from Sigma-Aldrich (St Louis, MO). Annealed oligos were quantified by BR ds DNA kit using Qubit spectrophotometer (Invitrogen, Carlsbad, CA). DNA fragments were separated by electrophoresis on 1% (w/v) or 2% (w/v) agarose gels, depending on the fragment size. PCR products were purified using GenElute™ PCR Clean-Up Kit (Sigma-Aldrich), after restriction digestion of the PCR mixture with DpnI (Thermo Fisher Scientific) for removal of circular templates. When required, DNA fragments were excised from gel and purified using Zymoclean™ Gel DNA Recovery Kit (Zymo Research, Irvine, CA). Plasmids were isolated from *E. coli* cultures using Sigma GenElute™ Plasmid Miniprep kit (Sigma-Aldrich).

### Entry-vector plasmids construction

All plasmids used in this study are listed in Table 2.

**Table 2 –.**
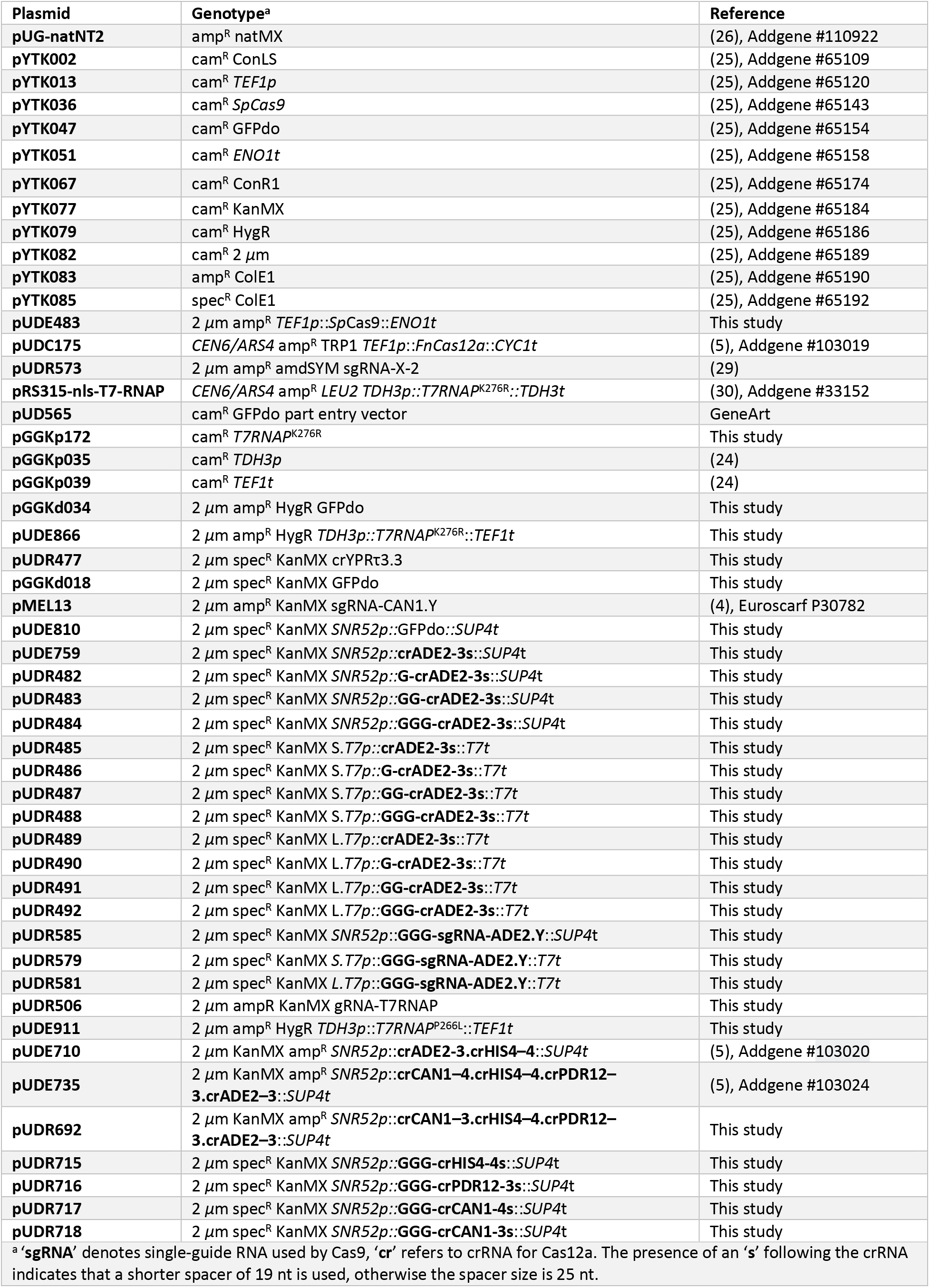
List of plasmids used in this study

The pUD565 plasmid (24), a GFP dropout (GFPdo) entry vector compatible with Yeast Toolkit parts (25), was ordered as synthetic gene from GeneArt (Thermo Fisher Scientific). GFPdo entry vectors for cloning of transcriptional unit were constructed following the BsaI Golden Gate reaction protocol described by Lee *et al*. (25). The GFPdo pGGKd018 plasmid was obtained by assembly of part plasmids pYTK002, pYTK047, pYTK067, pYTK077, pYTK082, pYTK085. The GFPdo pGGKd034 plasmid was constructed by assembly of part plasmids pYTK002, pYTK07, pYTK067, pYTK079, pYTK082, pYTK083. The GFPdo pUDE810, an entry vector for Cas12a-crRNAs, was constructed by Golden Gate assembly of pre-annealed primers 12647-12648 with the following PCR generated fragments: the pGGKd018 backbone with primers 12799-12800; the *SNR52* promoter amplified from the pMEL13 template (4) using primers 12645-13546; the GFPdo cassette bearing specific overhangs (GATC and ATCC) obtained by PCR amplification of primers 13547-12644 on pYTK047 (25).

### Construction of the dual Cas-expressing strain IMX1752

The construct for genomic integration of *cas9* gene consisted of a paired expression cassettes for introduction of *Streptococcus pyogenes cas9* nuclease and the *natNT2* marker into the *SGA1* locus (4). First, the *natNT2* marker was PCR amplified from pUG-natNT2 (Addgene plasmid #110922, (26)) using primers 10297 and 10298. This PCR product was cloned via Golden Gate together with pre-annealed primer pairs 10293-10294 and 10295-10296, and Yeast toolkit plasmids pYTK013, pYTK036, pYTK051, pYTK082, pYTK083 (25), resulting in plasmid pUDE483. The Cas9-natNT2 integration cassette was obtained by enzyme restriction of pUDE483 using EcoRI. The restriction mix was directly transformed into *S. cerevisiae* using the lithium acetate transformation protocol (27). Transformants were selected on YPD supplemented for nourseothricin. A single isolate, which was renamed IMX1714 (Table 1), was submitted to an additional transformation for the genomic integration of the Cas12a nuclease. For this, the sequence of *Francisella novicida cas12a* was amplified from pUDC175 (Addgene plasmid #103019, (5)) using primers 13553-13554. The obtained PCR product, carrying 60bp homology flanks to the X-2 integration site (28), was transformed in IMX1714 as previously described in (27), together with plasmid pUDR573 (29) for Cas9-mediated targeting at this genomic site. Transformants were selected on SM-Ac plates. Correct genomic integrations were confirmed by diagnostic PCR using primers listed in Table S1. After removal of the gRNA expression plasmid, the dual Cas9/Cas12a *S. cerevisiae* strain was stocked as IMX1752.

### Construction of the T7RNAP^K276R^-expressing strain IMX1905

First, the T7RNAP^K276R^ sequence was PCR amplified from plasmid pRS315-nls-T7-RNAP (Addgene plasmid #33152) (30) using primers 13543 and 13544, and the obtained PCR fragment stably cloned into entry vector pUD565, resulting in part plasmid pGGKp172. The T7RNAP^K276R^ transcriptional unit was assembled by Golden Gate cloning into plasmid pGGKd034, together with part plasmids pGGKp035 (*TDH3p*) and pGGKp039 (*TEF1t*) (24), leading to plasmid pUDE866. For genomic integration of the T7RNAP^K276R^, the previously characterized YPRCτ3 site of *S. cerevisiae* genome was chosen as recipient locus (31). Thus, a gRNA for Cas12a-mediated editing at this site was designed accordingly to guidelines provided in Swiat *et al*. (5). The gRNA for integration in YPRCτ3 was ordered as oligos 14142-14143 containing specific overhangs for Golden Gate assembly (GATC and ATCC). Oligo annealing and cloning into pUDE810 plasmid resulted in the crRNA-expressing plasmid pUDR477. Amplification of the T7RNAP^K276R^ integrative cassette was carried out on pUDE866 plasmid using primers 14022 and 14023, which contain repair ends of 60 bp homologous to the YPRCτ3 locus. This generated PCR fragments were co-transformed with plasmid pUDR477 into IMX1752 cells, as previously described (27). Yeast cells were selected on solid YPD plates supplemented with G418. Diagnostic PCR was performed on a single colony isolate, plasmid was recycled and the constructed strain was renamed IMX1905.

### Construction of T7RNAP mutants (IMX2030, IMX2031, IMX2032) and T7RNAP-overexpressing strains (IME459, IME475)

In order to alter the T7RNAP protein sequence, the *T7RNAP* gene of IMX1905 was *in vivo* mutated by means of the CRISPR-Cas9 editing machinery. A single gRNA was chosen for targeting the sequence surrounding DNA encoding amino acids at positions 266 and 276 (corresponding to 276 and 286 if considering the NLS) (Figure S3). For this, oligo 14284 was Gibson assembled by bridging to the pMEL13 (4) backbone, which was previously PCR amplified using primers 6005-6006. The obtained plasmid was renamed pUDR506. Repair oligos (Table S1) consist of 120-bp surrounding the T7RNAP targeted sequence with SNPs for P266L and/or R276K mutations, and carrying a silent mutation at the PAM sequence to avoid reiterative cutting. Plasmid pUDR506 and each double-stranded repair oligos were co-transformed into competent IMX1905 cells (27). Transformants were plated on YPD agar supplemented with G418. Screening of eight selected colonies was performed by SNP genotyping with primers listed in Table S1, following previously described procedures for SNP scoring (32). After SNPs validation and Sanger sequencing of the mutated T7RNAP sequence (Figure S3), strains were stocked as follow: IMX2030 (T7RNAP^P266L, K276R^) was renamed after the P266L amino acid substitution; IMX2031 (*wt*T7RNAP) expresses the wild-type T7RNAP, where the arginine at position 276 is changed into the native lysine; IMX2032 (T7RNAP^P266L^) resulted from simultaneous mutations of proline and arginine at positions 266 and 277 for the respective amino acid change in leucine and lysine.

For the T7RNAP^K276R^ overexpression, the dual Cas-expressing strain IMX1752 was transformed with plasmid pUDE866, following standard practice (27). Transformants were selected on YPD plates supplemented with hygromycin B and the strain was renamed as IME459. In parallel, transformation of IMX1752 with the empty vector pGGKd034 leaded to the control strain IME460. To overexpress the T7RNAP^P266L^ variant, the gene sequence of strain IMX2032 was PCR amplified from its isolated genomic DNA, using primers 10753 and 10768. The obtained PCR product was cloned by Golden Gate assembly into the episomal entry plasmid pGGKd034. The obtained plasmid, renamed pUDE911, was therefore transformed into IMX1752, transformants plated on selective YPD hygromycin B media and selected colonies stocked as IME475.

### Construction of gRNA expression cassettes

The gRNA cassettes for evaluation of *ADE2* deletion efficiencies mediated by Cas12a or Cas9 nucleases were prepared using the highly-efficient *ADE2-3* (5) or the *ADE2.y* (1) gRNAs, respectively. Each gRNA cassette was expressed from high-copy plasmid and comprised the gRNA sequence left-flanked by the RNAPolIII-dependent *SNR52p*, the minimal *T7p* ‘TAATACGACTCACTATA’ (*S.T7p*) or an extended *T7p* ‘GCCGGGAATTTAATACGACTCACTATA’ (*L.T7p*), with respective terminator sequences at the right flank. For the Cas12a-mediated targeting of other genes, previously characterized gRNAs were expressed as single gRNA-expressing cassette or as in an array-like arrangement: *HIS4* (HIS4-4), *PDR12* (PDR12-3) or *CAN1* (CAN1-4 or CAN1-3) (5).

All single gRNA-expressing plasmids were assembled by Gibson assembly reaction using the NEBuilder^®^ HiFi DNA Assembly Master Mix (New England Biolabs). Depending of the plasmid features, the backbone of the pUDE810 plasmid was amplified using different primer couples for specific homology overhangs. The backbone for assembly of Cas12a-gRNAs with *SNR52p/SUP4t* flanks was obtained by PCR amplification with primers 12710-5793. For T7RNAP-mediated expression of gRNAs via Cas12a, plasmid backbone was obtained by PCR amplification using either primers 14274-13713 (*S.T7p/T7t*) or 14275-13713 (*L.T7p/T7t*). Plasmid pUDE759 was assembled by *SNR52p/SUP4t* backbone fragment with annealed oligos 12713-12714. Single oligos were used for the Gibson assembly of Cas12a-gRNAs cassettes by single-stranded DNA bridging to each individual PCR-originated backbone fragments: the *SNR52p/SUP4t* derivatives pUDR482 (primer 14282), pUDR483 (primer 13750), pUDR484 (primer 14283), pUDR715 (primer 17328), pUDR716 (primer 17329), pUDR717 (primer 17330), pUDR718 (primer 17331); the *S.T7p/T7t*-related plasmids pUDR485 (primer 14280), pUDR486 (primer 13751), pUDR487 (primer 14281), pUDR488 (primer 13572); the *L.T7p/T7t* cognate plasmids pUDR489 (primer 14276), pUDR490 (primer 14277), pUDR491 (primer 14278) and pUDR492 (primer 14279).

For assembly of Cas9-gRNAs under *SNR52p*, the amplified pUDE810 backbone with *SNR52p*/*SUP4t* edges was mixed with annealed oligos 15508-15509 and the single-stranded oligo 14426 in a Gibson reaction, resulting in plasmid pUDR585. For Cas9-gRNAs expressed by T7RNAP, T7-edged plasmid backbones were PCR amplified using primers 14274-15287 (*S.T7p/sgRNA-T7t*) or 14275-15287 (*L.T7p/sgRNA-T7t*), generating a dsDNA fragment that additionally contains a partial sequence of the gRNA scaffold for Cas9. Gibson assemblies of annealed oligos 15290-15291 to either the *S.T7p/sgRNA-T7t* or *S.T7p/sgRNA-T7t* PCR-generated backbones were performed to obtain plasmids pUDR579 and pUDR581, respectively.

The control array of gRNAs expressed by plasmid pUDR692 was ordered as synthetic gene from GeneArt (Thermo Fisher Scientific). *SNR52* promoter and gRNA design principles previously elucidated were used (5). The synthetic gRNA array was flanked by BsaI sites and assembled by Golden Gate cloning into pGGKd018.

### Delivery of gRNA expression cassettes for single gene deletion in *S. cerevisiae*

Each gRNA expression cassette was transformed together with 1 μg of double-stranded deletion repair (Table S1) in exponentially growing *S. cerevisiae* cells (^~^2*10^7^ cells ml^−1^), accordingly to the lithium acetate transformation protocol (27). Genome editing via *in vitro* assembly described in Table 3 was prepared by transforming IMX1905 with 500 ng (^~^150 fmols) of each gRNA expression plasmid. For genome editing achieved via *in vivo* plasmid assembly, two linear PCR fragments were delivered with the transformation mix in IMX1905: i) 150 fmols of the specific gRNA cassette, systematically amplified from the respective *in vitro* constructed plasmid using primers 14584-14585; ii) 150 fmols of the linearized marked *2μ* backbone with 60 bp homology to each gRNA cassette, obtained from the amplification of pUDE810 with primers 11571-12378. To evaluate genome editing via delivery of linear gRNA expression cassette, each amplified gRNA cassette (150 fmols) was transformed with either 500 ng of circular pGGKd018 plasmid or with equimolar amount of two PCR fragments for the split plasmid selection using pGGKd018. These amplicons, having homologies for *in vivo* recircularization of the plasmid, were obtained by PCR amplification with primers 6815-9340 and primers 2398-12097.

**Table 3 –.**
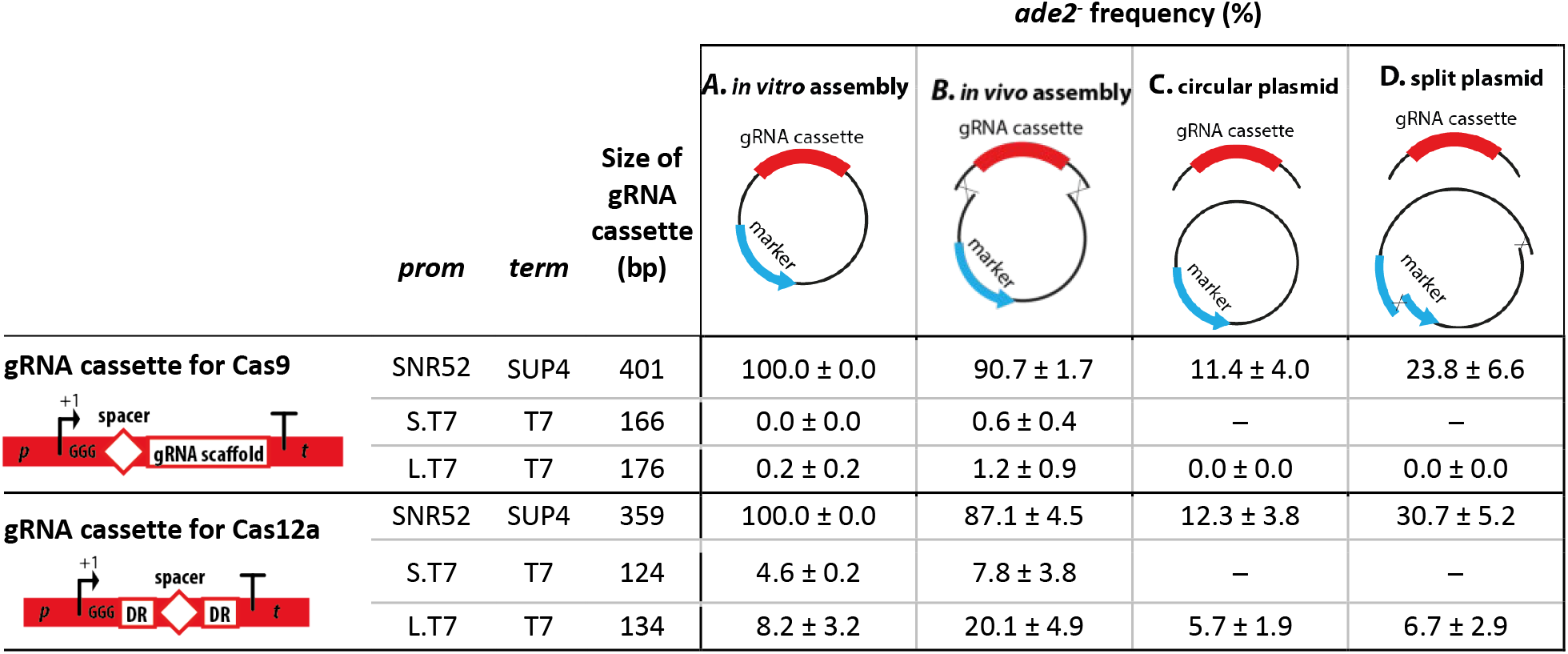
Comparing gRNA delivery methods. Cas9- and Cas12a-mediated DNA editing efficiency in IMX1905 (Table 1) transformed with different delivery methods for gRNA-expression cassette. A) *in vitro* pre-assembled plasmids; B) *in vivo* assembly after co-transformation of gRNA expression cassette and marker backbone with homology flanks; C) co-transformation of gRNA expression cassette with circular empty plasmid (pGGKd018); D) co-transformation of gRNA expression cassette with split empty plasmid (pGGKd018). The gRNA cassettes specific for Cas9 and Cas12a editing are depicted at the left of the table (*p*, promoter; DR, direct repeat; *t*, terminator) and their respective length in bp is reported. These were compared for expression under the RNAPolIII-dependent SNR52*p* (pUDR585 for Cas9, pUDR484 for Cas12a), the T7RNAP-dependent 17bp-long S.T7*p* (pUDR579 for Cas9, pUDR488 for Cas12a) or the T7RNAP-dependent 27bp-long L.T7*p* (pUDR581 for Cas9, pUDR492 for Cas12a). Editing efficiency is expressed as percentage of red colonies (*ade2^-^*). Data represent the average and standard deviation of biological triplicates.

All transformations were plated on selective YPD medium supplemented with G418. Efficiency of *ADE2* deletion is measured as number of red colonies on total CFU. For editing of other sites, diagnostic PCR was performed on a number of selected colonies using primer listed in Table S1.

### Preparation of gDNAs and genome editing via gEL DNA

Sequences of deletion repair fragments and gDNAs are listed as forward and reverse oligonucleotides in Table S1. Each forward and reverse oligo were mixed in equimolar amount, heated for 5 minutes at 95°C and cooled down to room temperature. As only exception, Cas9-mediated editing using gDNA with long T7 promoter was obtained by PCR amplification of two overlapping primers, the gRNA-specific forward for *ADE2.y* (16745) and the universal reverse carrying the Cas9-gRNA scaffold (16746). Concentrations of each double-stranded annealed oligos were measured for all pre-annealed oligos or the non-purified PCR-derived gDNA. 1 μg of each deletion repair and 4 μg of respective gDNA were mixed to 500 ng of split pGGKd18 plasmid for selection purposes and transformed into competent T7RNAP-expressing yeast cells accordingly to standard procedure (27). Transformants were selected on YPD plates supplemented with G418 if transforming a T7RNAP genomically integrated strains (IMX1905, IMX2030, IMX2031, IMX2032), or with G418 and hygromycin B if transforming a T7RNAP-overexpressing strains (IME459, IME475). Plasmid-base controls for multiplex via Cas12a gRNA-arrays were performed accordingly to Swiat *et al*. (5). Diagnostic PCR of selected colonies was done using primers listed in Table S1.

### Growth rate measurement

Strains were cultivated in 96-well plates containing SMD medium or SMD-urea supplemented with hygromycin B (30°C, 250 rpm). Growth was monitored by measuring optical density at 660 nm at regular time intervals using the Growth Profiler 960 (Enzyscreen B.V., Heemstede, The Netherlands). Maximum specific growth rates (*μ*_max_) were calculated using the equation 1: X=X_0_ e^μt^ in which μ indicates the exponential growth rate, from four independent biological cultures.

### Secondary structure prediction

The RNA secondary structure was predicted with the RNAstructure Web Server (https://rna.urmc.rochester.edu/RNAstructureWeb/) (33). Temperature was set to 30°C (303.15 K). Self-folding free energy are obtained via the same webtool.

## Results

### T7RNAP-expressing *S. cerevisiae* as a platform strain for Cas-mediated genome editing

For T7 RNA polymerase-(T7RNAP) based expression of gRNAs, the bacteriophage T7RNAP, previously functionally expressed in the yeast nucleus, was chosen (30). Flanked by the strong and constitutive *TDH3* promoter and *TEF1* terminator, T7RNAP was integrated in the genome of a *S. cerevisiae* strain from the CEN.PK family that constitutively expressed both Cas9 and Cas12a (strain IMX1752; Table 1). Sanger sequencing of the resulting strain IMX1905 revealed a missense mutation in the coding sequence of the T7RNAP as compared to the canonical sequence (https://www.uniprot.org/uniprot/P00573), which resulted in the replacement of a lysine by an arginine at the amino acid 276 of the polymerase (corresponding to amino acid 286 when considering the NLS) (Fig. S1). As the strain characterized by Dower and Rosbash (30) contained the same amino acid substitution and was proven to be functional in yeast, we decided to keep this variant (from now on referred to as T7RNAP^K276R^) to test the gEL DNA approach.

Physiological characterization revealed that IMX1905, co-expressing T7RNAP^K276R^, Cas9 and Cas12a, grew as fast as the prototrophic control strain CEN.PK113-7D in chemically defined medium supplemented with glucose as sole carbon source (specific growth rate of 0.42 ± 0.01 h^−1^ for IMX1905 and 0.45 ± 0.01 h^−1^ for CEN.PK113-7D) (Fig. 3). Expression of T7RNAP is therefore not toxic for *S. cerevisiae*.

**Figure 3:**
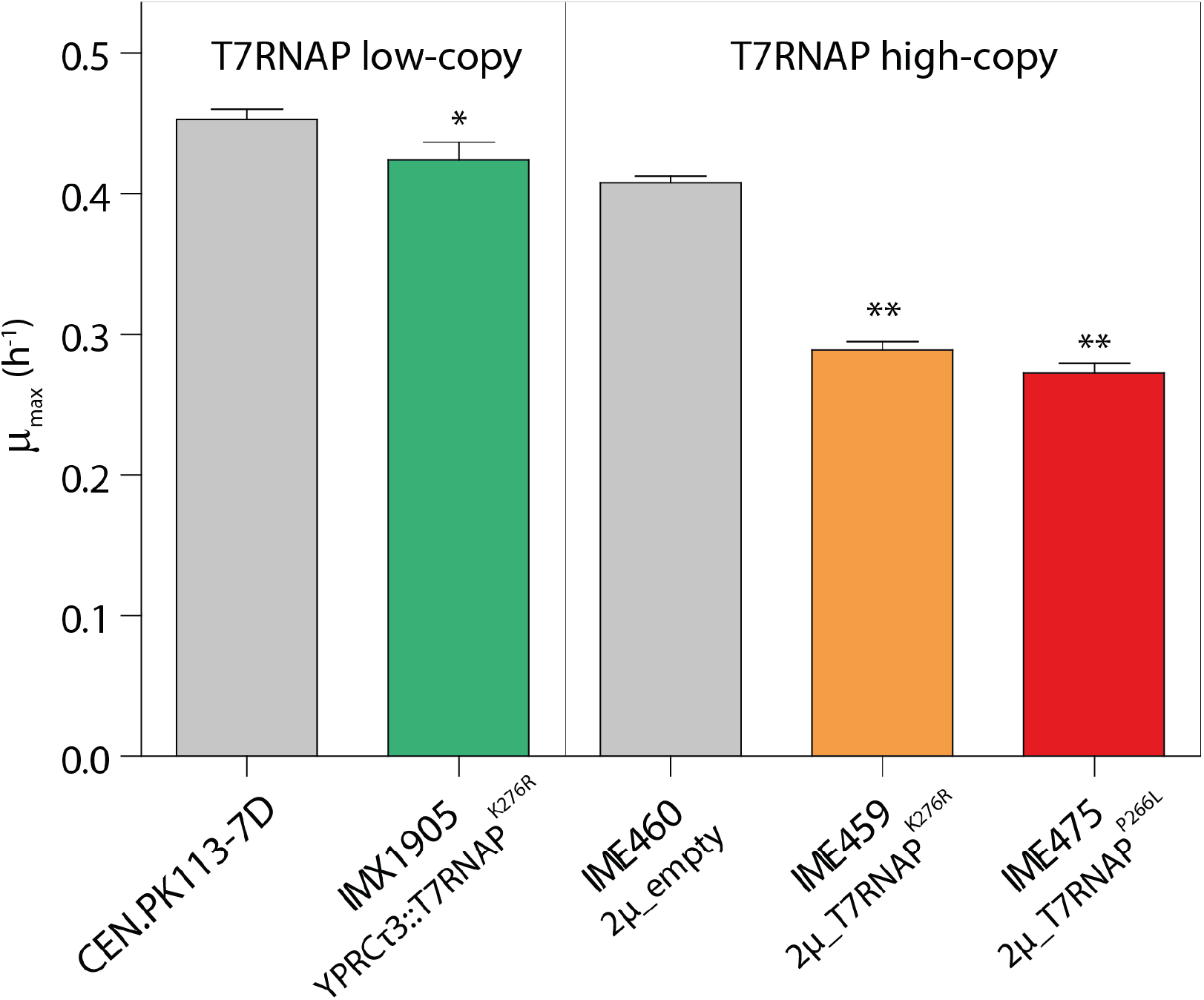
Physiological characterization of *S. cerevisiae* strains expressing T7RNAP. Maximum specific growth rates (*μ*_max_) of *S. cerevisiae* constitutively expressing T7RNAP^K276R^, Cas9 and Cas12a (IMX1905) and its control strain (CENPK.113-7D), or *S. cerevisiae* strains overexpressing T7RNAP^K276R^ (IME459) or T7RNAP^P266L^ (IME475) and its control strain carrying a 2μm multi-copy empty vector (IME460). Strains were cultivated in 96-well plate containing chemically defined medium supplemented with glucose as sole carbon source. Data points represent average and mean deviations of four biological replicates. *\*P* < 0.025, \*\**P* < 0.001, Student’s *t*-test was calculated compared to respective control strains CENPK.113-7D or IME460.

### T7RNAP enables gRNA expression from linear and circular DNA and promotes Cas9- and Cas12a-mediated DNA editing in *S. cerevisiae*

The activity of the T7RNAP^K276R^ in *S. cerevisiae* was evaluated by measuring the DNA editing efficiency of Cas9 and Cas12a guided by gRNAs expressed from T7 promoter. Two different T7 promoter lengths were tested, the minimal T7 promoter of 17 bp (TAATACGACTCACTATA; referred to as S.T7*p*), and an extended T7 promoter sequence of 27 bp (GCCGGGAATTTAATACGACTCACTATA; referred to as L.T7*p*) known to improve the stability of the T7RNAP-promoter complex *in vitro* (34). T7RNAP^K276R^-driven gRNA expression was compared with gRNA expression from the RNAPolIII-dependent *SNR52* promoter, largely adopted by the yeast community for CRISPR-Cas editing (1,5). Downstream all three promoters, at the initially transcribed region (ITS), a guanine triplet was added to increase the T7RNAP transcriptional activity (35). As previously reported for Cas9 (19), the addition of this guanine triplet strongly improved T7RNAP-mediated expression for Cas12a-based DNA editing (Fig. S2). Disruption of *ADE2*, leading to a red colony phenotype, was used to assess editing efficiency (36) in strain IMX1905 (Table 1). Spacers previously shown to guide Cas9 and Cas12a to *ADE2* with high efficiency were chosen (1,5). Cas9 and Cas12a have different requirements for functionality, which results to different compositions and size of the DNA cassette encoding the gRNA (from now on called gRNA cassette) (Table 3). The *ADE2.y* gRNA for Cas9 was used in its standard chimeric form including the trans-acting RNA (tracrRNA) (1). The *ADE2-3* gRNA for Cas12a was reduced to the minimal 19 nt-long spacer enclosed by the matured direct repeats (DR), as recently described (37). Thanks to Cas12a minimal requirements for DNA targeting and editing (no tracrRNA, small DR and spacer), the gRNA cassette for Cas12a were substantially smaller than those for Cas9 (Table 3).

The most popular method for gRNA delivery is via *in vitro* assembly of the gRNA expression cassette on a plasmid, and transformation of this circular plasmid to yeast. As expected, expression of the gRNA cassette from *SNR52p* using this delivery method led to high efficiency in editing of the *ADE2* gene in all colonies tested with both Cas9 and Cas12a (Table 3A). Conversely, T7RNAP^K276R^-based gRNA expression resulted in extremely low editing efficiency with Cas12a (4.6% ± 0.2% for the short and 8.2% ± 3.2% for the long T7 promoter) and null or negligible editing with Cas9 (Table 3A). Next to delivering a ready-made gRNA plasmid, two parts, one carrying the gRNA and the other the selection marker, were transformed into yeast. These two parts were flanked by 60 bp homologous sequences to enable *in vivo* circularization upon transformation. This delivery method enabled the transient availability of the gRNA cassette as linear DNA fragment. While reducing editing efficiency for SNR52*p*-based gRNA expression by ca. 10%, this method substantially increased *ADE2* editing by T7RNAP^K276R^-mediated expression of gRNAs for both Cas9 and Cas12a (Table 3B). Editing by Cas9 remained extremely low (around 1%), while up to 20% of the colonies displayed the disruption of *ADE2* by Cas12a (Table 3B). Next, to test whether the gRNA could be solely expressed from a linear DNA molecule, the gRNA cassette was delivered as double-stranded DNA fragment. A plasmid carrying a selectable marker was transformed in parallel for selection purposes. With this delivery method, editing efficiency with RNAPolIII-mediated gRNA expression was dramatically reduced to ca. 10% with both Cas9 and Cas12a (Table 3C). Editing efficiency for T7RNAP^K276R^-based gRNA expression was also reduced, but still detectable with ca 6% when using Cas12a. It has been shown that the efficiency of Cas9-mediated DNA editing can be increased by supplying a split plasmid during transformation, presumably by offering a selective advantage to cells that are proficient in homology directed repair (HDR) (9). Accordingly, a two-fold increase in *ADE2* editing efficiency was measured with both Cas9 and Cas12a when the gRNA were transcribed by RNAPolIII (Table 3C and D). However, using a circular or a split plasmid did not affect DNA editing for T7RNAP^K276R^-expressed gRNAs (Table 3C and D). The split marker approach combined with linear DNA delivery of the gRNA described in Table 3D was nonetheless kept for the following experiments.

Altogether these data demonstrate that gRNAs can be expressed from circular and linear DNA using the T7RNA^K276R^ polymerase in *S. cerevisiae*. Additionally, Cas12a leads to higher DNA editing efficiency than Cas9 when guided by T7RNAP^K276R^-expressed gRNAs. In all experiments, the long T7 promoter consistently led to two-to three-fold higher editing efficiencies than the short T7 promoter, suggesting that L.T7*p* drives higher gRNA expression (Table 3A and B). While delivery of gRNAs in the form of short linear DNA fragments enabled DNA editing, the observed editing efficiencies were low and required further optimization to turn the gEL DNA approach into an attractive and competitive DNA editing technique.

### Improving the efficiency of the T7RNAP^K276R^–based gEL DNA technique by optimizing the gDNA design

Aiming for cloning-free genome editing, the gEL DNA technique relies on the simple utilization of customized double-stranded DNA oligos (referred to as gDNAs) for the *in vivo* T7RNAP^K276R^-mediated expression of gRNAs. To reduce synthesis costs and increase compatibility with high-throughput strain construction, the size of the gDNA should be as small as possible and should not exceed 120 nt, the standard size limit of commercial, custom-made oligomers. In this respect, Cas12a presents a clear advantage as its gRNAs consists of a smaller structural part (DR) than the one required for DNA targeting by Cas9 (gRNA scaffold). Consequently, the small size of Cas12a gRNAs gives more flexibility in gDNA design regarding length of T7 transcriptional elements and presence of terminal DR or T7 terminator. Conversely, a minimal gDNA configuration for Cas9-mediated editing containing the S.T7p and the chimeric gRNA is 119-nt long (Fig. 4), which does not leave room for additional, potentially useful parts such as a longer T7 promoter or a T7 terminator. As previously, a triplet of guanine was appended to the T7 promoter for all tested gDNA.

**Figure 4:**
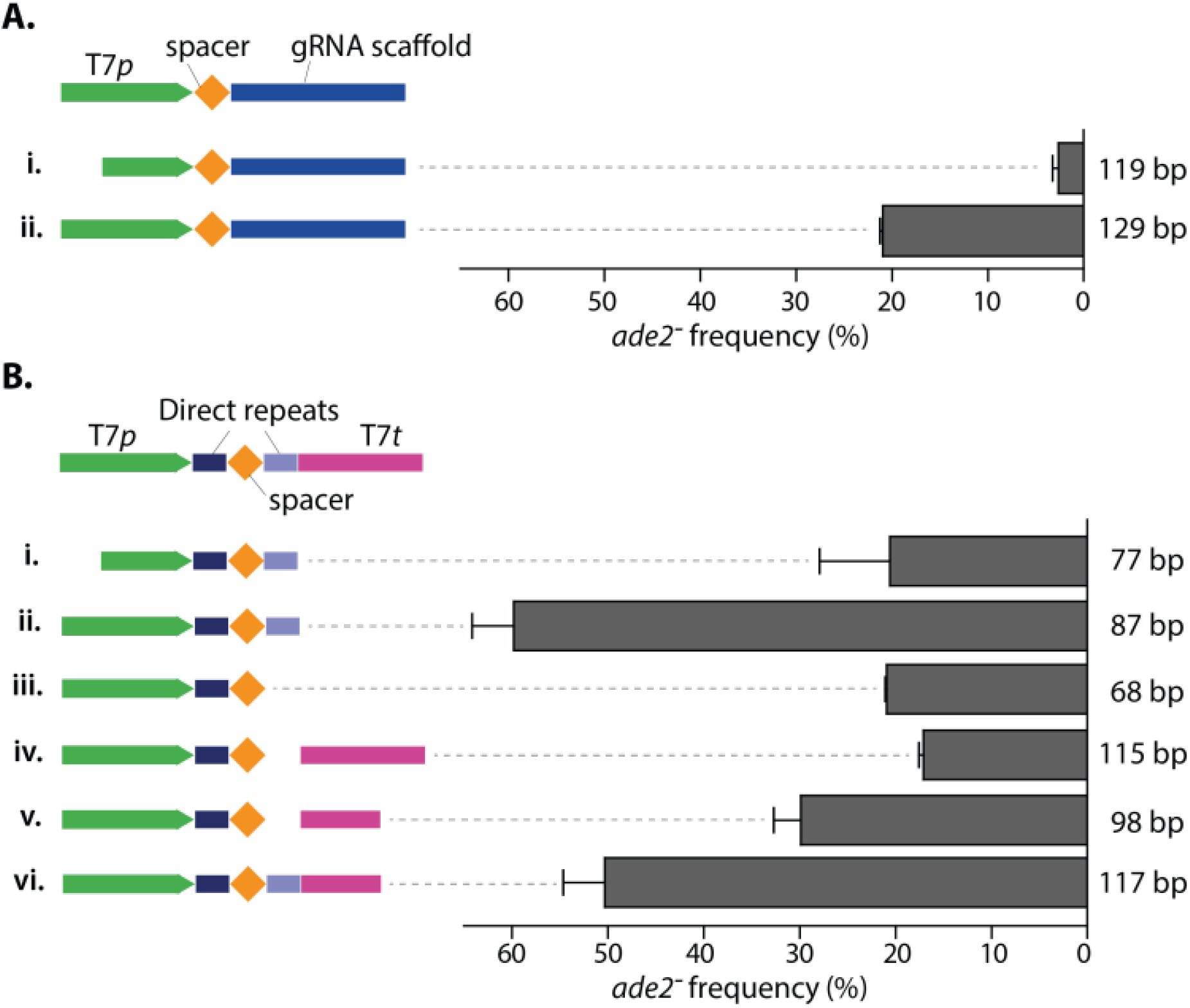
Optimization of Cas9 and Cas12a gDNA design. Editing efficiency of *ADE2* in strain IMX1905 transformed with gDNAs for cloning-free, T7RNAP-driven expression of gRNA. A) gDNA configurations for Cas9-mediated genome editing and respective editing efficiencies. B) gDNA configurations for Cas12a-mediated genome editing and their respective editing efficiencies. The size of each gDNA is specified on the right of the respective graph bar. Editing efficiency is expressed as percentage of red colonies (*ade2*) over the total number of colonies. Values represent the average and standard deviations of data obtained from three independent biological replicates.

For Cas9, two gDNA configurations were tested, one with the short and one with the long T7 promoter, followed by the *ADE2.y* spacer and the gRNA scaffold (Fig. 4A). As compared to the design presented in Table 3, no T7 terminator was added at the end of the gDNA. While the pre-annealed S.T7p gDNA can be directly transformed into IMX1905, the longer L.T7p gDNA (129 bp) had to be obtained by a preliminary PCR reaction using two primers with overlapping homologies (see Material and Methods section). Both gEL DNA configurations did enable Cas9-mediated DNA editing, marginal for the S.T7p (2.6% editing efficiency, Fig. 4A-i) but substantial for the L.T7*p* (21% editing efficiency, Fig. 4A-ii).

Six different gDNA configurations for the *ADE2-3* target were tested for Cas12a-mediated editing, differing in the size of T7 promoter and terminator as well as in the addition of a terminal DR and a T7 terminator (Fig. 4B). After simple pre-annealing of two complementary oligos *in vitro*, each gDNA was directly transformed into strain IMX1905. These data revealed that the terminal DR is important for efficient editing of *ADE2* irrespective to the presence of the T7 terminator, and that the presence of a T7 terminator is not required (Fig. 4B). They also further confirmed that the long version of the T7 promoter markedly increased DNA editing efficiency (Fig. 4B, i-ii). This design optimization enabled to increase the DNA editing efficiency to 60%, relying on the very simple transformation of yeast with a 87 nt-long oligonucleotide. This simple and efficient design, represented in Fig. 4B-ii, was implemented for the rest of the work with Cas12a.

The editing efficiencies shown in Fig. 4 were substantially higher for both Cas9 and Cas12a than those reported in Table 3D, in which a similar approach with linear gDNA delivery together with a split plasmid was used. This increased efficiency most probably resulted from the higher amount of gDNA used for the experiments presented in Fig. 4 (300 to 400-fold higher), which suggested that the abundance of delivered gDNA might be a key element for efficient DNA editing using gEL DNA.

### Improving the efficiency of the gEL DNA technique by optimizing sequence and expression levels of the T7RNAP

To further explore whether gDNA, and consequently gRNA availability might be limiting editing efficiency, gDNA transcription efficiency by the T7RNAP was explored. To this end, three additional T7RNAP variants were tested. All three variants were constructed from IMX1905, by inserting point mutations in the T7RNAP^K276R^ gene. In the first variant the K276R mutation was reverted into the wild-type T7RNAP (*wt*T7RNAP, strain IMX2031). The second variant carried the P266L mutation, known to reduce abortive transcription *in vitro* (38) (T7RNAP^P266L^, IMX2032) and the third variant carried the two mutations (T7RNAP^P266L^, ^R276K^, strain IMX2030). When tested with Cas12a, all four variants enabled DNA editing with a significantly higher efficiency for T7RNAP^P266L^ (Fig. 5), while T7RNAP^K276R^ and T7RNAP^P266L, R276K^ showed the lowest DNA editing efficiency (Fig. 5), suggesting that the K276R substitution is deleterious for T7RNAP transcription efficiency.

**Figure 5:**
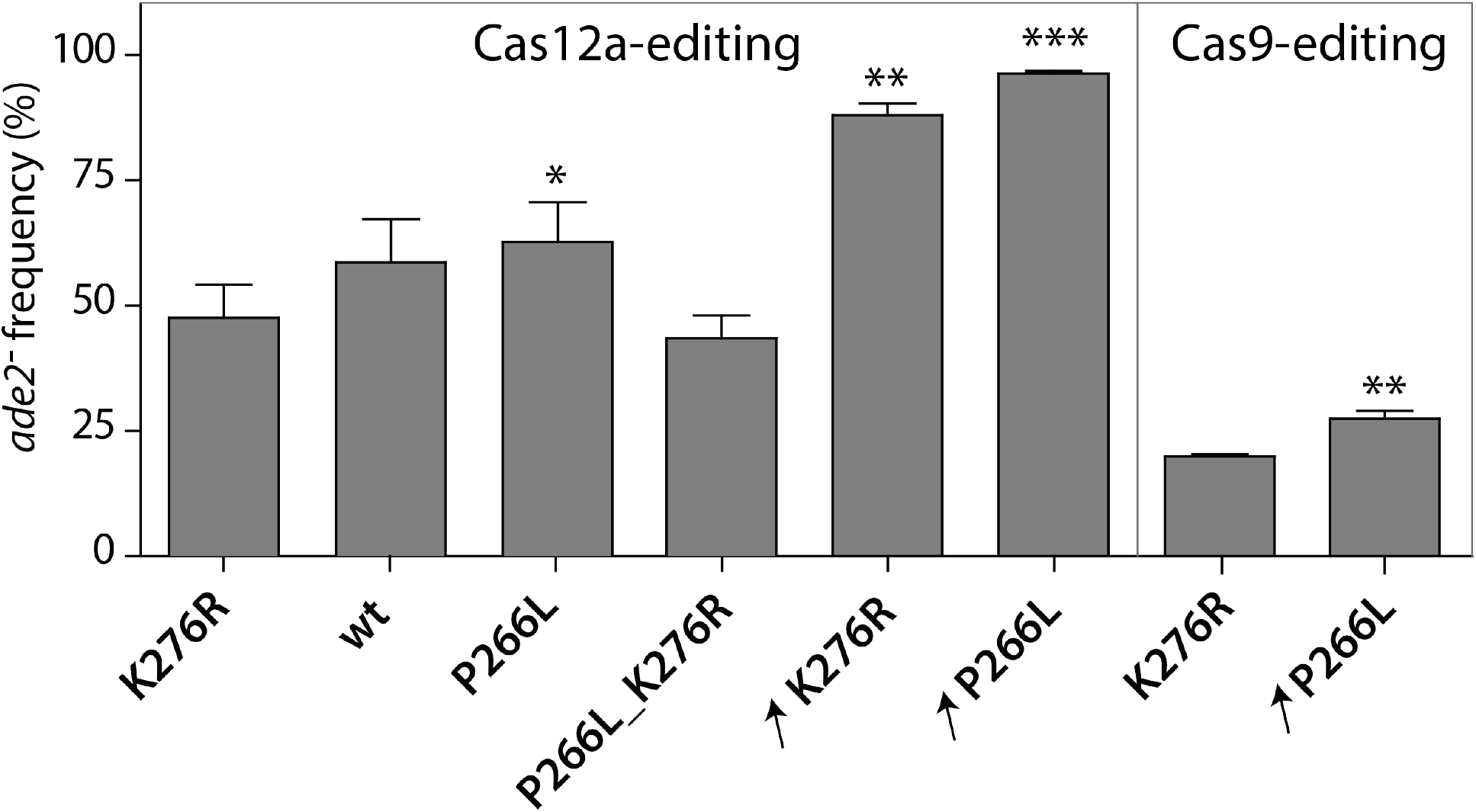
Comparison of Cas9 and Cas12a editing efficiency with T7RNAP variants. Efficiency of *ADE2* editing by Cas12- or Cas9-mediated gEL DNA in T7RNAP mutant or overexpression strains: IMX1905 (K276R); IMX2031 (wild-type, *wt*); IMX2032 (P266L); IMX2030 (P266L_K276R); IME459 (K276R overexpression, ↗K276R); IME475 (P266L overexpression, P266L). For Cas12a, transformed gDNA corresponds to annealed 15093-15094 oligos. For Cas9, transformed gDNA was obtained by PCR-derived fragment using overlapping primers 16745-16746. Editing efficiency is expressed as percentage of red colonies (*ade2^−^*). Values represent the average and standard deviations of data obtained from independent biological duplicates. \**P* < 0.05, \*\**P* < 0.025, \*\*\**P* < 0.001, Student’s *t*-test was calculated compared to respective control strain IMX1905 (K276R).

To further enhance gDNA transcription efficiency, the expression level of the T7RNAP was increased. The strains IME459 and IME475 were constructed by transformation with episomal plasmids harbouring the T7RNAP^R276K^ and the T7RNAP^P266L^ variants, respectively. While expression of T7RNAP from a single, integrated gene copy did not affect growth of *S. cerevisiae* (Fig. 3), expression from episomal vectors significantly reduced the growth rate when compared to a control strain carrying an empty episomal vector (0.29 ± 0.01 h^−1^ for IME459, 0.27 ± 0.01 h^−1^ for IME475 and 0.41 ± 0.00 h^−1^ for the control strain IME460, Fig. 3; Table 1). Overexpressing of either T7RNAP strongly increased the DNA editing efficiency by Cas12a, approaching 100% when using T7RNAP^P266L^ (Fig. 5). Overexpression of T7RNAP^P266L^ also increased DNA editing efficiency by Cas9, as compared to a single copy of T7RNAP^K276R^, but to a lesser extent (increase by 10%, Fig. 5). The gEL DNA approach remained much more efficient with Cas12a than with Cas9 (maximum efficiencies of 96% and 29%, respectively, Fig. 5)

Altogether these results revealed that the expression levels of the T7RNAP and consequently gRNA availability, play a key role for successful DNA editing by Cas12a in the gEL DNA system.

### gEL DNA enables Cas12a-mediated multiplex genome editing in *S. cerevisiae*

To test for multiplex genome editing, four gRNAs targeting *CAN1, HIS4, PDR12* and *ADE2*, previously shown to lead to 100% DNA editing efficiency by Cas12a when expressed from a RNAPolIII promoter, were selected ((5), Fig. 6A). As done for the *ADE2-3* target used for singleplex gEL DNA, these four additional gRNAs were shortened to a 19 bp-long spacer as compared to the previously described plasmid-based constructs (5). Oligos carrying the gDNA design shown in Fig. 4B-ii were ordered for each gRNA (Table S1) and transformed in duplex or quadruplex to IME475 overexpressing the T7RNAP^P266L^. Duplex targeting of *ADE2* and *HIS4* revealed that a vast majority of tested clones were edited (14 out of 16) and that 63% of the clones carried a double deletion (Fig. 6B). Out of the clones with single editing, none carried a single *HIS4* deletion, while duplex editing with *ADE2* was clearly a frequent event (Fig. S4). Quadruplex targeting resulted in a substantial fraction of clones without any editing (34%, Fig. 6C). The fraction of clones with a single editing event was very similar for duplex and quadruplex editing (25 and 30%, respectively). 23% and 13% of the clones carried double and triple editing, respectively and quadruplex editing was not observed (Fig. 6C). Remarkably, none of the tested clones displayed editing in *CAN1* (Fig. S5), suggesting that the *CAN1-4* gRNA failed to guide Cas12a to the targeted site. This lack of targeting might be explained by the fact that the *CAN1-4* gRNA contained an additional guanine triplet and was six nucleotides shorter than the *CAN1-4* gRNA originally tested by Swiat and co-workers. To test this hypothesis, the *CAN1-4* gDNA (GGG at his 5’ and a 19 bp-long spacer) was expressed from a plasmid with the *SNR52* promoter and tested for editing efficiency. Out of eight selected colonies, none resulted in a *CAN1* deletion (Fig. S6), a complete loss in editing efficiency that is likely due to the disruption of the gRNA stem-loop structure (Fig. 6D). A new *CAN1* spacer with a predicted secondary structure displaying the gRNA stem-loop was therefore selected for *CAN1* targeting (*CAN1-3* (5), Fig. 6D). Expressed from a plasmid with the *SNR52* promoter, *CAN1-3* led to 100% *CAN1* editing with Cas12a (Fig. 6D). However, when tested for multiplexing using the gEL DNA, *CAN1-3* rarely led to editing of *CAN1* by Cas12a (Fig. 6E). A single *CAN1* editing event was observed out of 30 clones tested (Fig. S7) and, remarkably, this event was concomitant with the editing of the three other targets, leading to a single clone with quadruple DNA editing (Fig. 6E). In the quadruplex editing experiments with *CAN1-4* and *CAN1*-3, the fraction of clones with single, double and triple DNA editing was comparable (roughly 30, 25 and 10% respectively, Fig. 6).

**Figure 6:**
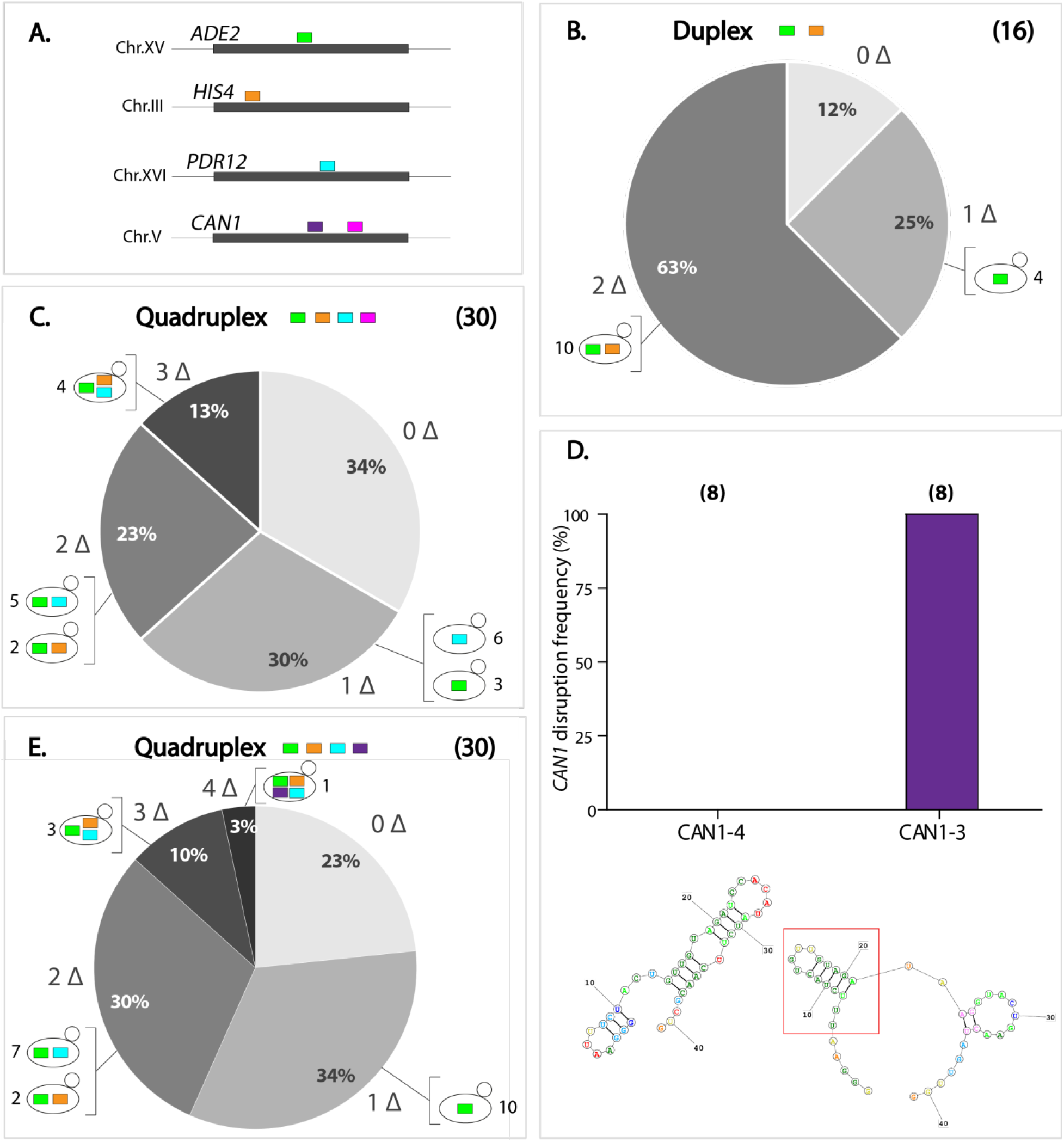
Multiplex genome editing by Cas12a-mediated using the gEL DNA approach. (A) Targeted sites for deletion of *ADE2* (ADE2-3, green), *HIS4* (HIS4-4, orange), *PDR12* (PDR12-3, cyan) and *CAN1* (CAN1-4, pink; CAN1-3, violet) genes. (B) Percentage of transformants obtained from double gDNAs delivery: ADE2-3 and HIS4-4. (C) Fraction of selected colonies upon transformation with four gDNAs: ADE2-3, HIS4-4, PDR12-3 and CAN1-4. (D) Verification of single editing efficiency of CAN1 targets expressed from plasmid pUDR717 (CAN1-4) or pUDR718 (CAN1-3), with prediction of gRNAs secondary structure. (E) Fraction of selected colonies upon transformation with four gDNAs: ADE2-3, HIS4-4, PDR12-3 and CAN1-3. Number of verified clones is indicated between brackets and diagnostic PCRs are reported in Supplementary Figures (Fig. S4, S5, S6). Zero (0Δ), single (1Δ), double (2Δ), triple (3Δ) or quadruple (4Δ) deletion are indicated at the outside ends of each fraction. Type of obtained deletions are specified with the respective colour of the target. Number of colonies are also stated next to each depiction. Prediction of the gRNA stem-loop for Cas12a recognition is highlighted by a red square.

Following the approach described by Swiat and coworkers, two cRNA arrays were tested for quadruplex genome editing. Both plasmids carried the *HIS4-4, ADE2-3* and *PDR12-3* gRNAs, but pUDE735 expressed *CAN1-4* (Fig. S5) while pUDR692 expressed *CAN1-3* (Fig. S7). As previously observed, the number of colonies obtained after transformation was extremely low (below ten colonies), as compared to the number of colonies obtained for quadruplex editing with the gEL DNA approach (over 150 colonies).

## Discussion

The future of the CRISPR-Cas-based genome editing heads towards the development of fast and low-cost methodologies for strain construction. The gEL DNA approach presented in this study expands the CRISPR-Cas genome editing toolbox of *S. cerevisiae* with an entirely cloning-free and very efficient strategy for single or double genetic modification in *S. cerevisiae*. By simply transforming pre-annealed 87-long, complementary DNA oligonucleotides into competent yeast cells, cost and time of strain construction can be reduced to their bare minimum. Any chosen gRNA cassette can be delivered independently or in combination with other gRNA cassettes, making this technique very versatile and highly suitable for high-throughput, combinatorial strain construction. Akin to other CRISPR-based techniques for genome engineering increasing the number of simultaneously targeted sites strongly affects the efficiency of multiplexed gEL DNA (Table S2, Fig. 6) (7). The results obtained in this study suggest that this efficiency can be further enhanced. For instance, increasing T7RNAP and gDNA abundance substantially increased singleplex gene editing (Fig. 4, Fig. 5), indicating that gRNA abundance is a key factor for efficient DNA editing. While the toxicity of plasmid-borne T7RNAP expression showed that its abundance cannot be further increased in *S. cerevisiae*, the efficiency of the gEL DNA could be further enhanced by T7RNAP protein engineering or by expression of DNA-dependent RNAP variants from other bacteriophages (*i.e*. T3, SP6 or K11) that are able to transcribe from short promoters and from linear DNA templates (39–41). Another aspect to consider is the stability of the gDNA. While other methods deliver gRNA in the form of plasmids that are very stable *in vivo*, the linear nature of the gDNA make it prone to degradation by native exonucleases. Further studies should explore the stability of gDNA and gRNA during transformation and test whether chemical stabilization of the linear gDNA (by phosphorotioate derivatives or 2’-ribose modification for instance (42,43)) enhances gRNA availability and thereby DNA editing. There are therefore several promising avenues to further improve multiplex DNA editing with the gEL DNA approach.

Out of the five gRNA tested in this study, one failed to guide Cas12a for gene editing. Remarkably, for this guide (*CAN1-4*) the folding prediction suggested the whole disruption of the direct repeat as a consequence of the 5’-addition of the guanine triplet, while the other four guides displayed typical gRNA secondary structures with the required stem-loop structure (Fig. 6 and Table S3) (44). In agreement with these observations, a recent study about the Cas12a-gRNA functionality suggests that the disruption of the direct repeat pseudoknot structure by pairing to the spacer sequence might lead to loss of gRNA targeting ability (37). Additionally, inhibition of the gRNA processing and consequently of Cas12a activity seems to be due to the positional effect of a stable secondary structure flanking the direct repeat (45). It has been recently advised that the terminator should be spaced-out by a 24 nt-long spacer to avoid steric effects with the pseudoknot formation and thereby allow correct gRNA folding (37). Our findings support these theories, since a gRNA flanked by the short, 30-nt T7 terminator sequence that lacks the stem-loop structure has a 1.8-fold higher *ADE2* editing efficiency than a gRNA with the longer, 47-nt T7 terminator (Fig. 4B, iv-v). Prediction of the gRNA structure is therefore essential to optimize Cas12a-based DNA editing with the gEL approach.

Despite efforts to improve editing with both Cas9 and Cas12a, the latter proved to be more efficient for DNA editing with the gEL DNA method. The causes for Cas9 lower efficiency remain to be elucidated, but the observation that increasing T7RNAP abundance hardly affects DNA editing by Cas9 (increased by 1.4%; Fig. 5) suggests that gRNA abundance is not the factor impairing Cas9 activity. While the length of the gDNA might be another obstacle for Cas9 implementation with the gEL DNA approach, it could be overcome by expressing the tracrRNA separately from the gRNA (3).

In conclusion, the gEL DNA methodology is not only an extremely valuable tool for genome editing in *S. cerevisiae*, but has a yet greater potential thanks to its portability to other organisms. Expression of gRNAs using the host machinery or *in vivo* burden of gRNA-expression plasmids can present serious obstacles for CRISPR/Cas9 based editing (2,46–49). By introducing a T7RNAP and gDNA oligos, the gEL DNA approach dissociates gRNA production from the host polymerase and from plasmid templates, thereby entirely removing these obstacles.

## Supplementary data

Supplementary Data are available at online.

## Acknowledgment

We thank Melanie Wijsman and Ewout Knibbe for constructing strains IMX1714 and IMX1752, respectively, and Sofia Dashko for cloning plasmid pGGKd034.

## Funding

This research received funding from the Netherland Organization of Scientific Research (NWO)

## Notes

### Competing Interest Statement

The authors have declared no competing interest.

