## Supplementary data for "gEL DNA, a cloning- and PCR-free method for CRISPR-based multiplexed genome editing"

### Supplementary Information

#### Figure S1 – Alignment of the T7RNAP from Dower and Rosbash with the wild type sequence

```
CLUSTAL O(1.2.4) multiple sequence alignment

Reference      ----------MNTINIAKNDFSDIELAAIPFNTLADHYGERLAREQLALEHESYEMGEAR 50
IMX1905        MLEPPKKKRKVNTINIAKNDFSDIELAAIPFNTLADHYGERLAREQLALEHESYEMGEAR 60
                         :*************************************************

Reference      FRKMFERQLKAGEVADNAAAKPLITTLLPKMIARINDWFEEVKAKRGKRPTAFQFLQEIK 110
IMX1905        FRKMFERQLKAGEVADNAAAKPLITTLLPKMIARINDWFEEVKAKRGKRPTAFQFLQEIK 120
               ************************************************************

Reference      PEAVAYITIKTTLACLTSADNTTVQAVASAIGRAIEDEARFGRIRDLEAKHFKKNVEEQL 170
IMX1905        PEAVAYITIKTTLACLTSADNTTVQAVASAIGRAIEDEARFGRIRDLEAKHFKKNVEEQL 180
               ************************************************************

Reference      NKRVGHVYKKAFMQVVEADMLSKGLLGGEAWSSWHKEDSIHVGVRCIEMLIESTGMVSLH 230
IMX1905        NKRVGHVYKKAFMQVVEADMLSKGLLGGEAWSSWHKEDSIHVGVRCIEMLIESTGMVSLH 240
               ************************************************************

Reference      RQNAGVVGQDSETIELAPEYAEAIATRAGALAGISPMFQPCVVPPKPWTGITGGGYWANG 290
IMX1905        RQNAGVVGQDSETIELAPEYAEAIATRAGALAGISPMFQPCVVPPRPWTGITGGGYWANG 300
               *********************************************:**************

Reference      RRPLALVRTHSKKALMRYEDVYMPEVYKAINIAQNTAWKINKKVLAVANVITKWKHCPVE 350
IMX1905        RRPLALVRTHSKKALMRYEDVYMPEVYKAINIAQNTAWKINKKVLAVANVITKWKHCPVE 360
               ************************************************************

Reference      DIPAIEREELPMKPEDIDMNPEALTAWKRAAAAVYRKDKARKSRRISLEFMLEQANKFAN 410
IMX1905        DIPAIEREELPMKPEDIDMNPEALTAWKRAAAAVYRKDKARKSRRISLEFMLEQANKFAN 420
               ************************************************************

Reference      HKAIWFPYNMDWRGRVYAVSMFNPQGNDMTKGLLTLAKGKPIGKEGYYWLKIHGANCAGV 470
IMX1905        HKAIWFPYNMDWRGRVYAVSMFNPQGNDMTKGLLTLAKGKPIGKEGYYWLKIHGANCAGV 480
               ************************************************************

Reference      DKVPFPERIKFIEENHENIMACAKSPLENTWWAEQDSPFCFLAFCFEYAGVQHHGLSYNC 530
IMX1905        DKVPFPERIKFIEENHENIMACAKSPLENTWWAEQDSPFCFLAFCFEYAGVQHHGLSYNC 540
               ************************************************************

Reference      SLPLAFDGSCSGIQHFSAMLRDEVGGRAVNLLPSETVQDIYGIVAKKVNEILQADAINGT 590
IMX1905        SLPLAFDGSCSGIQHFSAMLRDEVGGRAVNLLPSETVQDIYGIVAKKVNEILQADAINGT 600
               ************************************************************

Reference      DNEVVTVTDENTGEISEKVKLGTKALAGQWLAYGVTRSVTKRSVMTLAYGSKEFGFRQQV 650
IMX1905        DNEVVTVTDENTGEISEKVKLGTKALAGQWLAYGVTRSVTKRSVMTLAYGSKEFGFRQQV 660
               ************************************************************

Reference      LEDTIQPAIDSGKGLMFTQPNQAAGYMAKLIWESVSVTVVAAVEAMNWLKSAAKLLAAEV 710
IMX1905        LEDTIQPAIDSGKGLMFTQPNQAAGYMAKLIWESVSVTVVAAVEAMNWLKSAAKLLAAEV 720
               ************************************************************

Reference      KDKKTGEILRKRCAVHWVTPDGFPVWQEYKKPIQTRLNLMFLGQFRLQPTINTNKDSEID 770
IMX1905        KDKKTGEILRKRCAVHWVTPDGFPVWQEYKKPIQTRLNLMFLGQFRLQPTINTNKDSEID 780
               ************************************************************

Reference      AHKQESGIAPNFVHSQDGSHLRKTVVWAHEKYGIESFALIHDSFGTIPADAANLFKAVRE 830
IMX1905        AHKQESGIAPNFVHSQDGSHLRKTVVWAHEKYGIESFALIHDSFGTIPADAANLFKAVRE 840
               ************************************************************

Reference      TMVDTYESCDVLADFYDQFADQLHESQLDKMPALPAKGNLNLRDILESDFAFA    883
IMX1905        TMVDTYESCDVLADFYDQFADQLHESQLDKMPALPAKGNLNLRDILESDFAFA    893
               *****************************************************
```

Protein alignment of T7RNAP from IMX1905 (derived from Dower and Rosbash (1)) to reference sequence (https://www.uniprot.org/uniprot/P00573). The SV40 nuclear localization signal (NLS) is highlighted in yellow. Substitution of the lysine (K) at position 276 into an arginine (R) (aa 286 including the NLS) is highlighted in green.

#### Figure S2 – Impact of guanine addition to T7 promoters on *ADE2* editing efficiency

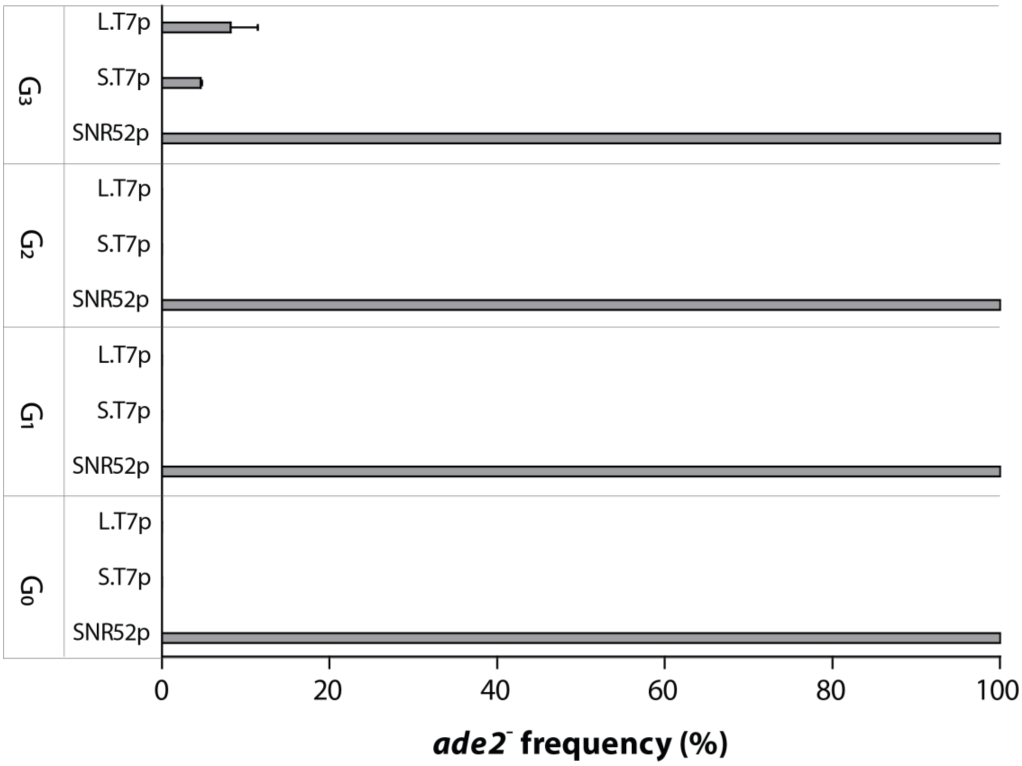


Cas12a-mediated DNA editing efficiency in IMX1905 (Table 1) transformed with plasmids for expression of *ADE2-3* gRNA under different promoters with increasing number of appended guanines ($G_x$ with x from 1 to 3) at the initially transcribed sequence (ITS). Editing efficiencies were compared for gRNAs expressed under RNAPolIII-dependent SNR52*p* (pUDE759, $G_0$; pUDR482, $G_1$; pUDR483, $G_2$; pUDR484, $G_3$), the T7RNAP-dependent 17bp-long S.T7*p* (pUDR485, $G_0$; pUDR486, $G_1$; pUDR487, $G_2$; pUDR488, $G_3$) or the T7RNAP-dependent 27bp-long L.T7*p* (pUDR489, $G_0$; pUDR490, $G_1$; pUDR491, $G_2$; pUDR492, $G_3$). These are expressed as percentage of red colonies (*ade2*$^-$). Data represent the average and standard deviations of biological duplicates.

#### Figure S3 - Diagnostic PCR and sequencing for the T7RNAP variants

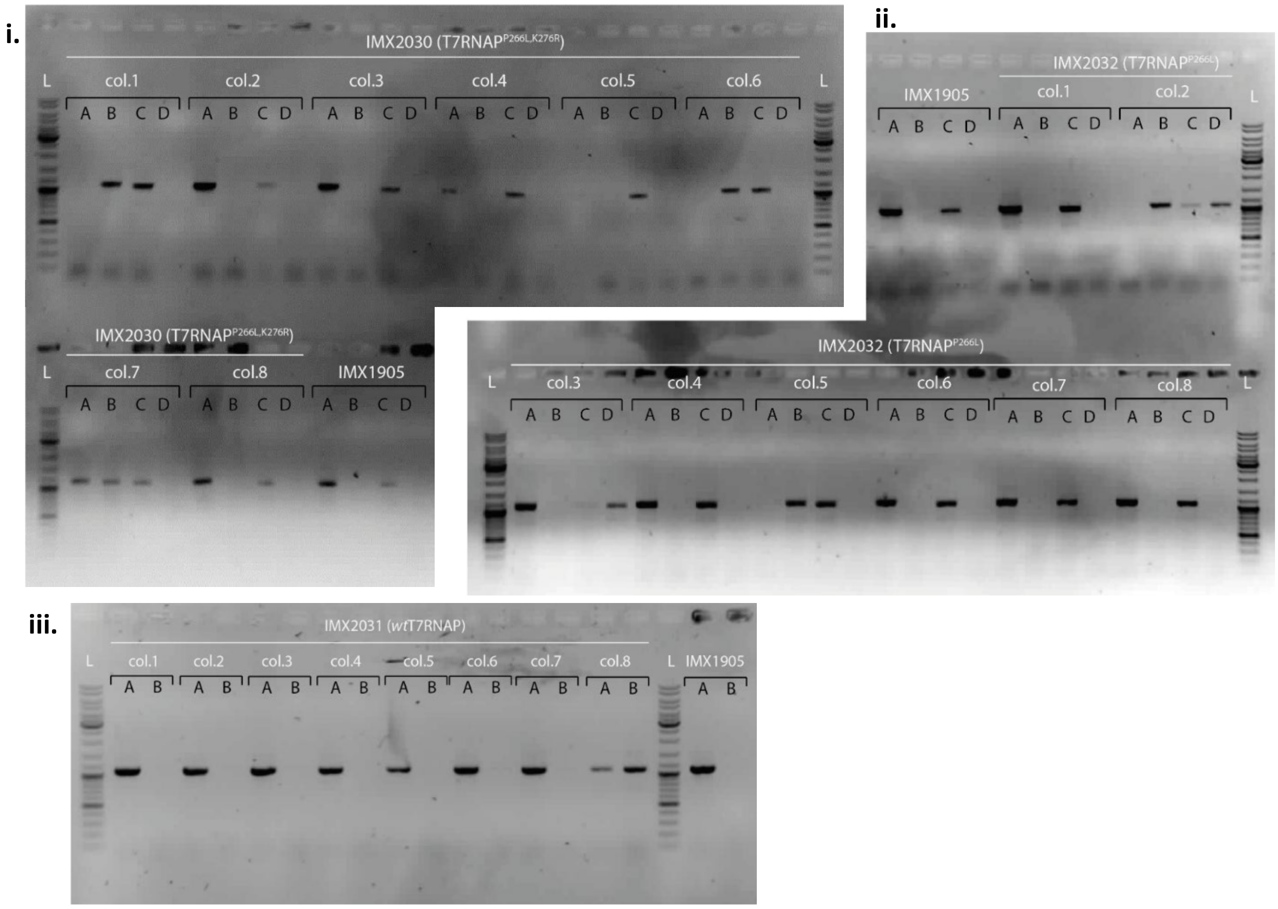


iv.

```
                 730         740         750         760         770         780         790         800         810
                   *           *           *           *           *           *           *       T   *           *
IMX1905   CGCCAAAATGCTGGCGTAGTAGGTCAAGACTCTGAGACTATCGAACTCGCACCTGAATACGCTGAGGCTATCGCAACCCGTGCAGGTGCG
IMX2030   .............................................................................A............
IMX2031   .............................................................................A............
IMX2032   .............................................................................A............
                                                                                       T

                 820         830         840         850         860         870         880         890         900
                   *       P   *           *           *       K   *           *           *           *           *
IMX1905   CTGGCTGGCATCTCTCCGATGTTCCAACCTTGCGTAGTTCCTCCTAGGCCGTGGACTGGCATTACTGGTGGTGGCTATTGGGCTAACGGT
IMX2030   ................T.........................................................................
IMX2031   ..............................................A...........................................
IMX2032   ................T.............................A...........................................
                          L                             R
```

SNP genotyping of colonies obtained by IMX1905 (Table 1) transformation with pUDR506 (Table 2) and respective repair fragments (Table S1) for the construction of strain IMX2030 (T7RNAP$^{P266L, K276R}$), IMX2031 (wild-type T7RNAP), IMX2032 (T7RNAP$^{P266L}$). i) ii) iii) Agarose gels with diagnostic PCR were performed on eight colonies picked from each transformation. Presence of bands A and B indicates positive colonies for IMX2030 (i), bands B and C for IMX2032 (ii) or bands A and B for IMX2031 (iii). iv) Sequencing data alignment of selected colonies highlight the silent mutation at the PAM sequence (in bold) (ACC→ACA) and the two mutations for the P266L (CCG→CTG) and K276R (AGG→AAG) amino acid substitutions (in green). Modified nucleotide triplets are framed, SNPs are highlighted in red. Targeted protospacer is underlined.

#### Figure S4 – Diagnostic PCR of duplex gEL DNA for *HIS4* and *ADE2* deletions

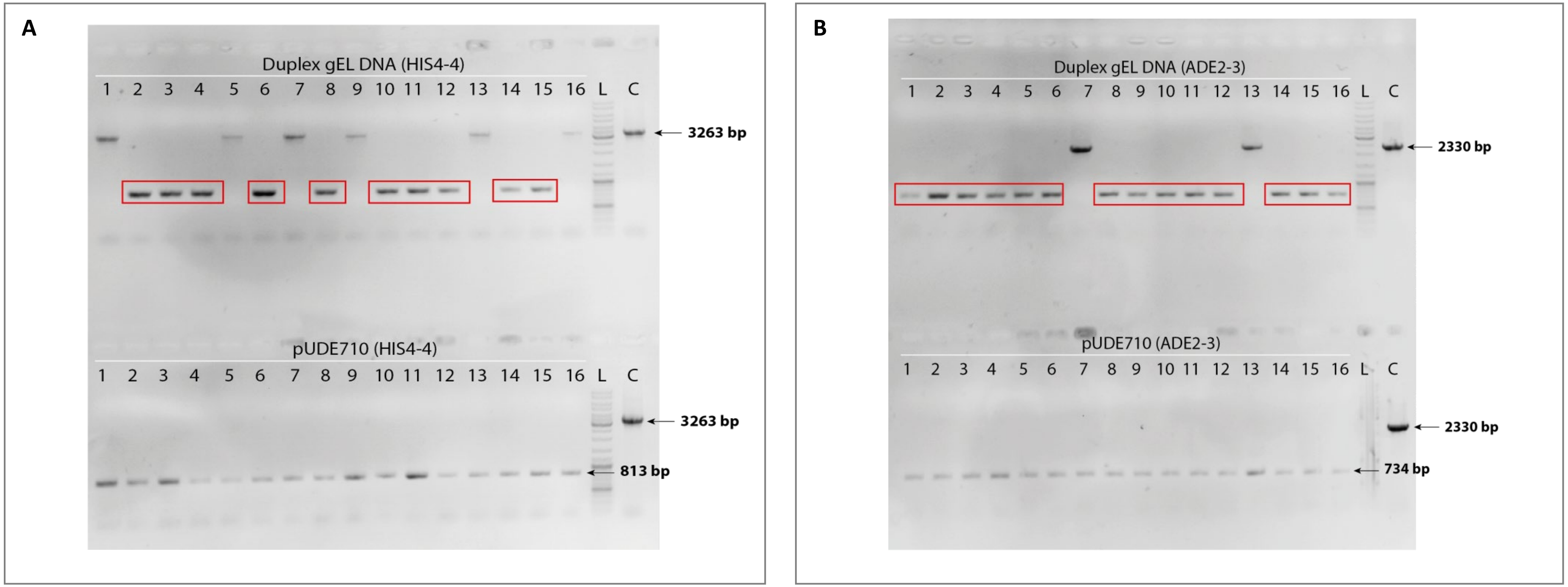


Diagnostic PCR of IME475 transformed with gDNAs for *HIS4* (GGG.crHIS4-4s) and *ADE2* (GGG.crADE2-3s) and respective repair fragments for gene deletions. Agarose gels with diagnostic PCR was performed on 16 picked colonies. Selected colonies from control array plasmids pUDE710 (crRNAs with 25 bp spacer) are also shown for each diagnostic gel. A: HIS4. B: ADE2. Expected fragment sizes are indicated and edited colonies are framed by a red square. L: ladder. C: negative control (CEN.PK113-7D).

Figure S5 - Diagnostic PCR for quadruplex gEL DNA of *HIS4, PDR12, CAN1* and *ADE2* deletions

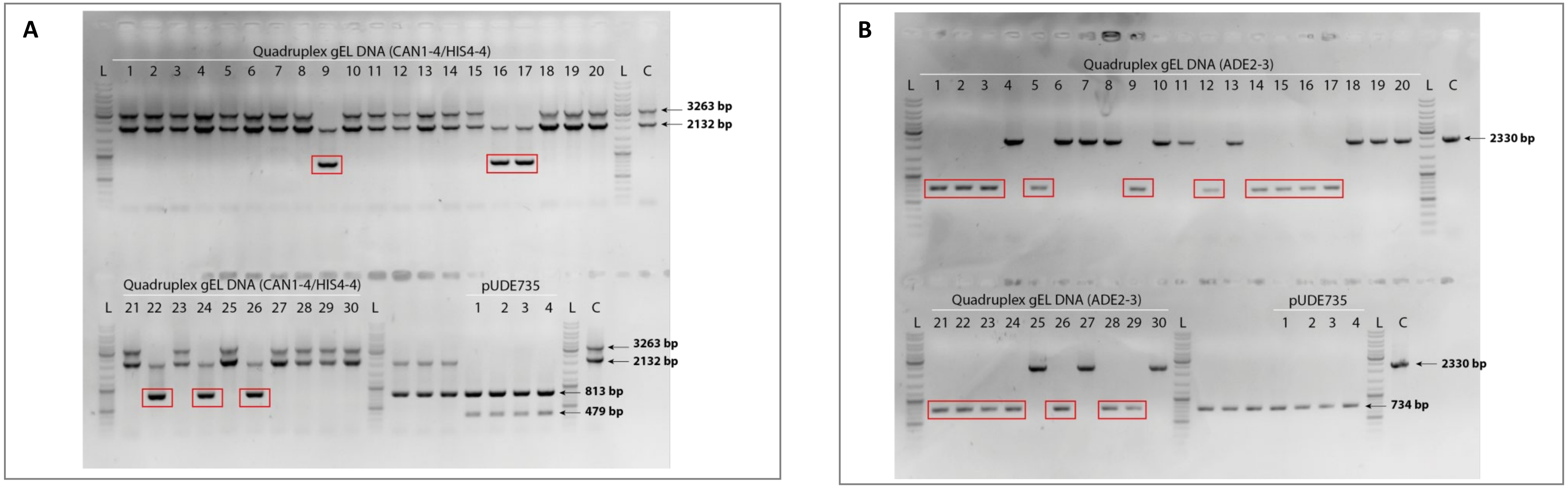

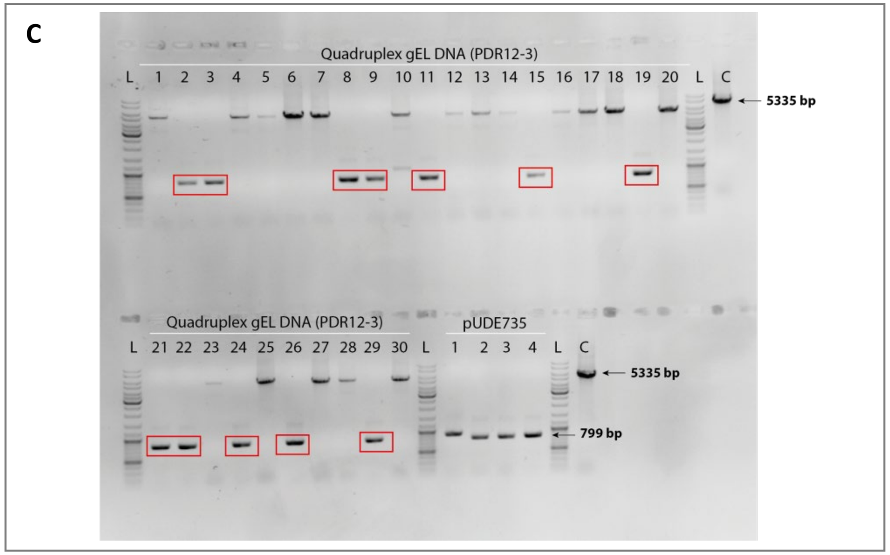


Diagnostic PCR of IME475 transformed with gDNAs for *CAN1* (GGG.crCAN1-4s), *HIS4* (GGGcrHIS4-4s), *ADE2* (GGG.crADE2-3s) and *PDR12* (GGG.crPDR12-3s) and respective repair fragments for gene deletions. Agarose gels with diagnostic PCR was performed on 30 picked colonies. Selected colonies from control array plasmid pUDE735 (crRNAs with 25 bp spacer) are also shown for each diagnostic gel. A: CAN1-HIS4. B: ADE2. C: PDR12. Expected fragment sizes are indicated and edited colonies are framed by a red square. L: ladder. C: negative control (CEN.PK113-7D).

Figure S6. - Diagnostic PCR for single targets *HIS4-4s, PDR12-3s, CAN1-4s, CAN1-3s*

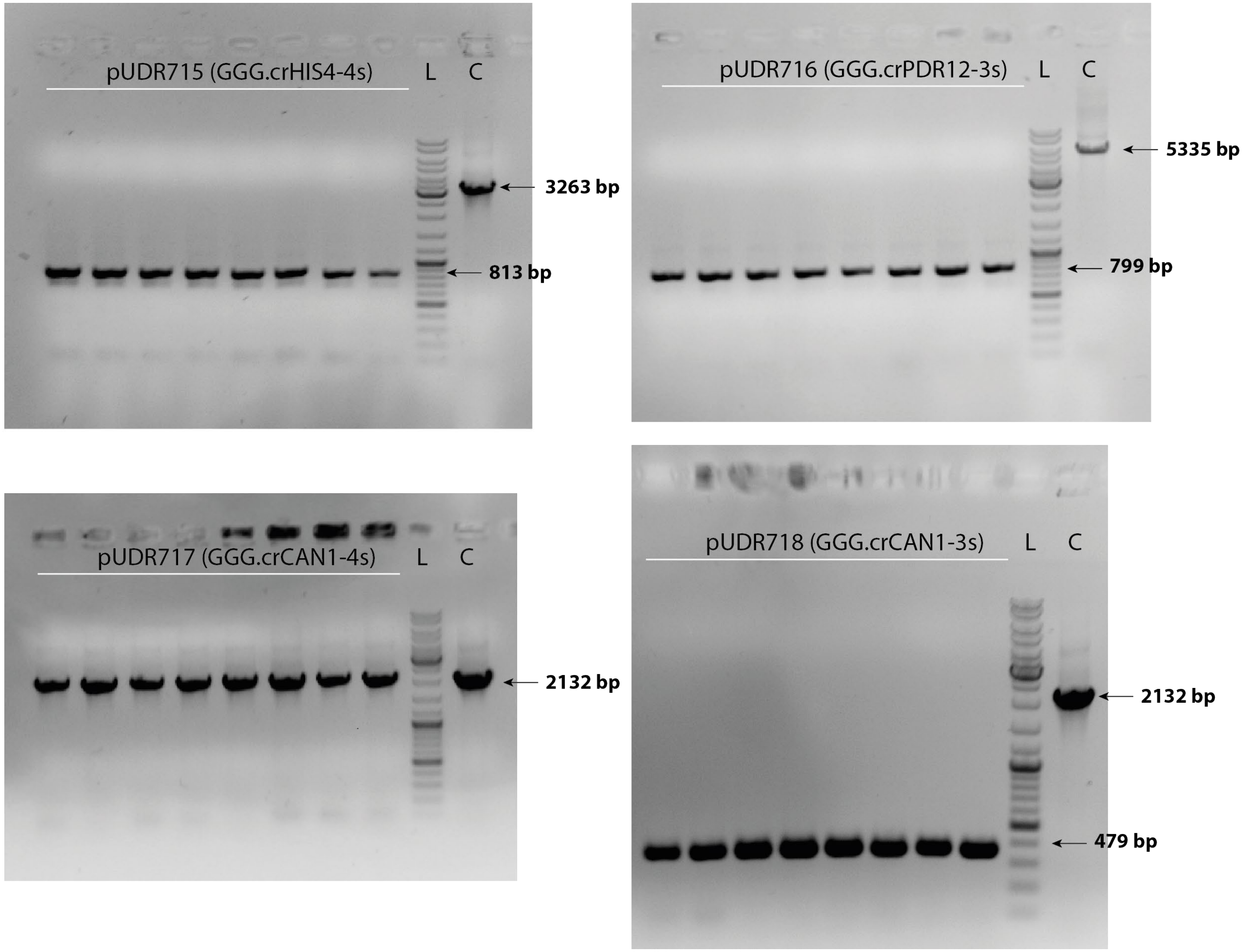


Diagnostic PCR of IME475 transformed with plasmids expressing GGG.crHIS4-4s (pUDR715), GGG.crPDR12-3 (pUDR716), GGG.crCAN1-4(pUDR717) or GGG.crCAN1-3 (pUDR718) and respective repair fragments for these gene deletions. Agarose gels with diagnostic PCR was performed on 8 selected colonies. Expected fragment sizes are indicated. L: ladder. C: negative control (CEN.PK113-7D).

Figure S7. - Diagnostic PCR for quadruplex gEL DNA of *HIS4, PDR12, CAN1* and *ADE2* deletions

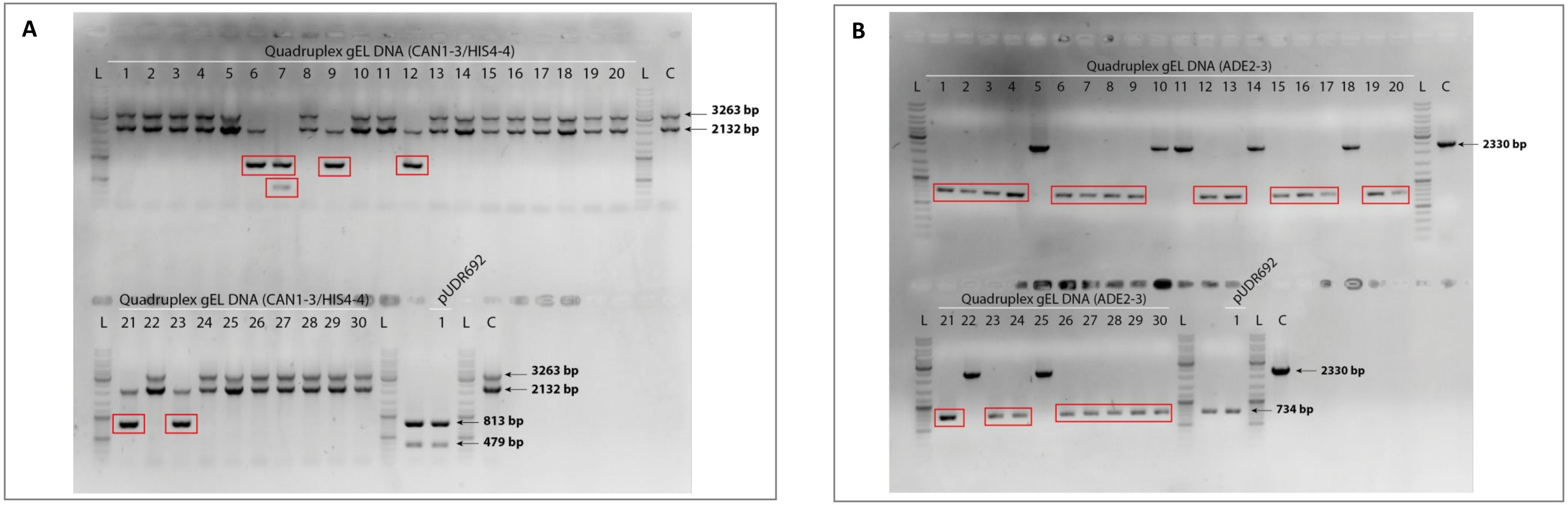

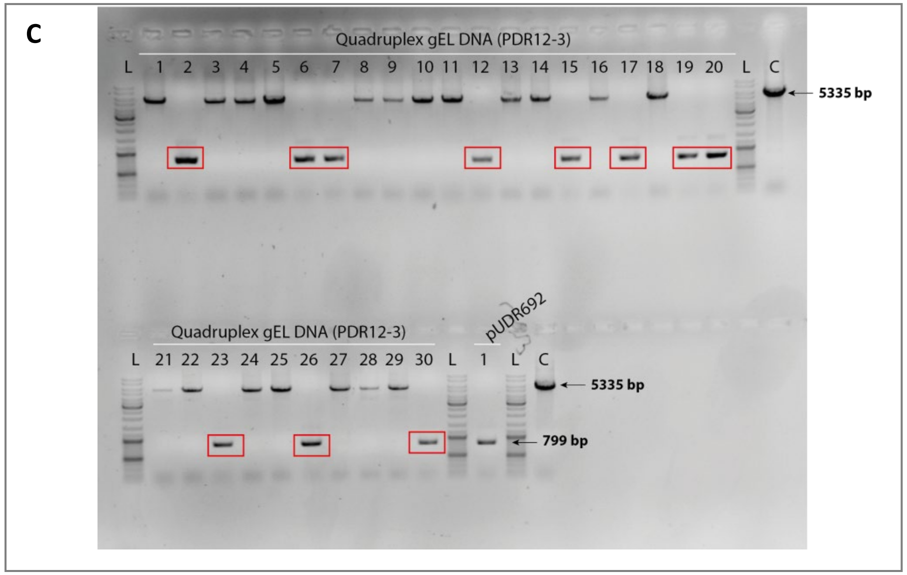


Diagnostic PCR of IME475 transformed with gDNAs for *CAN1* (GGG.crCAN1-3s), *HIS4* (GGG.crHIS4-4s), *ADE2* (GGG.crADE2-3s) and *PDR12* (GGG.crPDR12-3s) and respective repair fragments for gene deletions. Agarose gels with diagnostic PCR was performed on 30 picked colonies. Selected colonies from control array plasmid pUDR692 (crRNAs with 25 bp spacer) are also shown for each diagnostic gel. A: CAN1-HIS4. B: ADE2. C: PDR12. Expected fragment sizes are indicated and edited colonies are framed by a red square. L: ladder. C: negative control (CEN.PK113-7D).

#### Table S1- Primers table

List of primers used in this study.

| Number | Primer name | Sequence (5’ -> 3’) | Purpose |
|---|---|---|---|
| **Construction and genotyping of Cas9/Cas12a-expressing *S. cerevisiae* strain (IMX1752)** | | | |
| **10293** | YTK_1_sga1F fw | GGTCTCACCCTGAATTCTATCTGTTCTGTTACACAAGAAATCGTACATTTACAATATAGTGATAATCGTGGACTAGAGCAAGATTTCAAATAAGTAACAGCAGCAAAAACGAGAGACC | Construction of pUDE483 |
| **10294** | YTK_1_sga1F rv | GGTCTCTCGTTTTTGCTGCTGTTACTTATTTGAAATCTTGCTCTAGTCCACGATTATCACTATATTGTAAATGTACGATTTCTTGTGTAACAGAACAGATAGAATTCAGGGTGAGACC | |
| **10295** | YTK_6_sga1F fw | GGTCTCATACAAGCGAGAAGTATACACAAGTGTATTTCCTAGATATTTACATCAAATATATATATACTTATTTACAAAACTCTGATATTATAAATTAATTAGAATTCGAGTAGAGACC | |
| **10296** | YTK_6_sga1F rv | GGTCTCTACTCGAATTCTAATTAATTTATAATATCAGAGTTTTGTAAATAAGTATATATATATTTGATGTAAATATCTAGGAAATACACTTGTGTATACTTCTCGCTTGTATGAGACC | |
| **10297** | pUG-NAT YTK fw | GGTCTCAGCTGCCAGCTGAAGCTTCGTAC | |
| **10298** | pUG-NAT YTK rv | GGTCTCATGTAGCATAGGCCACTAGTGGATC | |
| **13553** | pTEF1_X2_integration_FW | GCTGAAGATTTATCATACTATTCCTCCGCTCGTTTCTTTTTTCAGTGAGGTGTGTCGTGAAAGCTGGAGCTCATAGCTTC | Cas12a cassette amplification |
| **13554** | tCYC1_X2_integration_RV | ATTCTCGCCAAGGCATTACCATCCCATGTAAGAACGGAATAAAACAGCATTCGAAGGTTAGCCGCAAATTAAAGCCTTCG | |
| **4666** | c PEPC fw | CTCTTTCCTGTAGGTCAGGTTG | Diagnostic *Sp*Cas9 in *sga1* locus |
| **7468** | Q_OutsideSGA_FW | GGCTCTGGATCCGTTATCTG | |
| **7479** | Q_OutsideSGA_RV | GGACGTTCCGACATAGTATC | |
| **8410** | 1000bp band_seq_2_rev | CGACGAAGAAAAAGAAACGAGG | |
| **7376** | x-2_outside | GGTCTAGGCCTGCATAATCG | Diagnostic *Fn*Cas12a in X-2 locus |
| **10245** | checkFnCpf1_sgaI_Fw | TGGCTGATGTCAACGGAAAC | |
| **10246** | checkFnCpf1_sgaI_Rv | CTGATGCACACGGAGCTCAG | |
| **10888** | X2 outside rev | ACCGACCCTAACAAACTCTC | |
| **Construction of Cas12a-gRNA entry vector plasmid (pUDE810)** | | | |
| **12645** | fw_pSNR52 | GGGGGTCTCAAACGTCTTTGAAAAGATAATGTATGATTATGCTTTC | Amplification of *SNR52p* |
| **13546** | pSNR52_rv | GGGGGTCTCAGATCATTTATCTTTCACTGCGGAGA | |
| **13547** | GFPdo_fw | GGGGATCTGAGACCGAAAGTGAAACG | Amplification of GFP dropout |
| **12644** | rv_GFPdropout | GGGGGATTGAGACCTATAAACGCAGAAAGG | |

| | | | |
|---|---|---|---|
| **12647** | fw_DR-SUP4t | GGGGGTCTCAATCCAATTTCTACTGTTGTAGATTTTTTTTGTTTTTTATGTCTGCTGTGA GACCCCC | Assembly of *FnCpf1* direct repeat and *SUP4t* |
| **12648** | rv_DR-SUP4t | GGGGGTCTCACAGCAGACATAAAAAACAAAAAAAATCTACAACAGTAGAAATTGGAT TGAGACCCCC | |
| **12799** | BB_fw | GGGGGTCTCAGCTGCCAATGAGAC | Amplification of pGGKd018 backbone |
| **12800** | BB_rv | GGGGGTCTCaCGTTCGTCTGCAATTATC | |
| **<u>Construction and genotyping of T7 RNA polymerase in *S. cerevisiae* strains (IMX1905)</u>** | | | |
| **13543** | YTK_T7_fw | GCATCGTCTCATCGGTCTCATATGCTCGAGCCTCCAAAAAAGAAGAGA | Construction of pGGKp172 |
| **13544** | YTK_T7_rv | ATGCCGTCTCAGGTCTCAGGATTTACGCGAACGCGAAGTCCGA | |
| **3416** | I-rev | CTGCGTTATCCCCTGATTCTGTGG | Diagnostic of pUDE866 or pUDE911 |
| **9895** | pGAL check RV | GCGAGGCACATCTGCGTTTC | |
| **14022** | YPRCtau3_ConS_fw | AGAATGATTTACAATCTAGTCGCAAAAACAAGTACAGTGCTGACGTCCCATCTTTTAAT GCATGCGCGGATGACACGAACTCA | T7RNAP cassette amplification |
| **14023** | YPRCtau3_Con1_rv | CATTACCAATGAATGCTGTTTTTGCAGAAATAACGAGATATCTGCAATAAAAGCAAAA GTCCTTGATCTGTCGGGTGTCTGCT | |
| **14142** | gRNA-YPRCtau3-3_fw | GGGGGTCTCAGATCAATTTCTACTGTTGTAGATGTGTGAAGGTACCTATTGCATCCAGA GACCGGG | Construction of pUDR477 |
| **14143** | gRNA-YPRCtau3-3_rv | CCCGGTCTCTGGATGCAATAGGTACCTTCACACATCTACAACAGTAGAAATTGATCTGA GACCCCC | |
| **13262** | YPRCtau3 dg RV | GCCTCCCCTAGCTGAACAAC | Diagnostic T7RNAP into YPRCτ3 and Sanger sequencing |
| **13710** | T7in_rv | GAGTAACACCGTAAGCCAGC | |
| **10216** | Ttef1_sc fw | GGAGATTGATAAGACTTTTCTAGTTG | |
| **13261** | YPRCtau3 dg FW | AATACGAGGCGAATGTCTAGG | |
| **<u>Construction of plasmids for *ADE2* editing via Cas9 or Cas12a nuclease</u>** | | | |
| **12710** | fw_pHiMex | TTGTTTTTTATGTCTGCTGCCAAT | Backbone amplification for *in vitro* plasmid construction (RNAPolIII) |
| **5793** | pCAS9 rv | GATCATTTATCTTTCACTGCGGAG | |
| **13713** | T7t-Con_fw | TAGCATAACCCCTTGGGGCCTCTAAACGGGTCTTGAGGGGTTTTTTGGGGCCTTTCTGC GTTTATAGGT | Backbone amplification for *in vitro* plasmid construction (*T7t*) |
| **14274** | T7p-Con_rv | TATAGTGAGTCGTATTAGTATTGCGACGAATTGCCACG | Backbone amplification for *in vitro* plasmid construction (S.*T7p*) |
| **14275** | longT7p-Con_rv | TATAGTGAGTCGTATTAAATTCCCGGCGTATTGCGACGAATTGCCACG | Backbone amplification for *in vitro* plasmid construction (L.*T7p*) |
| **15287** | Cas9gRNA_T7t_BB_fw | *GGCACCGAGTCGGTGGTGC*TAGCATAACCCCTTGGGGCCTCTAAACGGGTCTTGAGG GGTTTTTTGTTTTTTTGTTTTTTATGTCTGCTGCCA | Backbone amplification for *in vitro* plasmid construction (Cas9-gRNA scaffold, *T7t*) |

| | | | |
|---|---|---|---|
| **14426** | SNR52pGGG_gADE2.Y | GCATGTTTCGGCGTTCGAAACTTCTCCGCAGTGAAAGATAAATGATCGGGACTTGAAGATTCTTTAGTGTGTTTTAGAGCTAGAAATAGCAAGTTAAAATAAGGCTAGTCCGTTATCAAC | Construction of pUDR585 |
| **15508** | struct_SUP4t_fw | GTTTTAGAGCTAGAAATAGCAAGTTAAAATAAGGCTAGTCCGTTATCAACTTGAAAAAGTGGCACCGAGTCGGTGGTGCTTGTTTTTTATGTCTGCTGCCAAT | |
| **15509** | struct_SUP4t_rv | ATTGGCAGCAGACATAAAAAACAAGCACCACCGACTCGGTGCCACTTTTTCAAGTTGATAACGGACTAGCCTTATTTTAACTTGCTATTTCTAGCTCTAAAAC | |
| **15290** | T7p-GGGgADE2y_fw | TAATACGACTCACTATAGGGACTTGAAGATTCTTTAGTGTGTTTTAGAGCTAGAAATAGCAAGTTAAAATAAGGCTAGTCCGTTATCAACTTGAAAAAGTGGCACCGAGTCGGTGGTGC | Construction of pUDR579 or pUDR581 |
| **15291** | T7p-GGGgADE2y_rv | GCACCACCGACTCGGTGCCACTTTTTCAAGTTGATAACGGACTAGCCTTATTTTAACTTGCTATTTCTAGCTCTAAAACACACTAAAGAATCTTCAAGTCCCTATAGTGAGTCGTATTA | |
| **12713** | crADE2-3s_fw | GCAGTGAAAGATAAATGATCAATTTCTACTGTTGTAGATCCGGTTGTGGTATATTTGGAATTTCTACTGTTGTAGATTTTTTTTGTTTTTTATGTCTGCTGCCAAT | Construction of pUDE759 |
| **12714** | crADE2-3s_rv | ATTGGCAGCAGACATAAAAAACAAAAAAAATCTACAACAGTAGAAATTCCAAATATACCACAACCGGATCTACAACAGTAGAAATTGATCATTTATCTTTCACTGC | |
| **14282** | gADE2-3sG | AAACTTCTCCGCAGTGAAAGATAAATGATCGAATTTCTACTGTTGTAGATCCGGTTGTGGTATATTTGGAATTTCTACTGTTGTAGATTTTTTTTGTTTTTTATGTCTGCTGCCAATG | Construction of pUDR482 |
| **13750** | gADE2-3sGG | AAACTTCTCCGCAGTGAAAGATAAATGATCGGAATTTCTACTGTTGTAGATCCGGTTGTGGTATATTTGGAATTTCTACTGTTGTAGATTTTTTTTGTTTTTTATGTCTGCTGCCAATG | Construction of pUDR483 |
| **14283** | gADE2-3sGGG | AAACTTCTCCGCAGTGAAAGATAAATGATCGGGAATTTCTACTGTTGTAGATCCGGTTGTGGTATATTTGGAATTTCTACTGTTGTAGATTTTTTTTGTTTTTTATGTCTGCTGCCAATG | Construction of pUDR484 |
| **14280** | T7p_gADE2-3 | GTCGCAATACTAATACGACTCACTATAAATTTCTACTGTTGTAGATCCGGTTGTGGTATATTTGGAATTTCTACTGTTGTAGATTAGCATAACCCCTTGGGGCCTCTAAACGGG | Construction of pUDR485 |
| **13751** | T7pG_gADE2-3 | GTCGCAATACTAATACGACTCACTATAGAATTTCTACTGTTGTAGATCCGGTTGTGGTATATTTGGAATTTCTACTGTTGTAGATTAGCATAACCCCTTGGGGCCTCTAAACGGG | Construction of pUDR486 |
| **14281** | T7pGG_gADE2-3 | GTCGCAATACTAATACGACTCACTATAGGAATTTCTACTGTTGTAGATCCGGTTGTGGTATATTTGGAATTTCTACTGTTGTAGATTAGCATAACCCCTTGGGGCCTCTAAACGGG | Construction of pUDR487 |
| **13752** | T7pGGG_gADE2-3 | GTCGCAATACTAATACGACTCACTATAGGGAATTTCTACTGTTGTAGATCCGGTTGTGGTATATTTGGAATTTCTACTGTTGTAGATTAGCATAACCCCTTGGGGCCTCTAAACGGG | Construction of pUDR488 |
| **14276** | longT7p_gADE2-3 | GCCGGGAATTTAATACGACTCACTATAAATTTCTACTGTTGTAGATCCGGTTGTGGTATATTTGGAATTTCTACTGTTGTAGATTAGCATAACCCCTTGGGGCCTCTAAACGGG | Construction of pUDR489 |
| **14277** | longT7pG_gADE2-3 | GCCGGGAATTTAATACGACTCACTATAGAATTTCTACTGTTGTAGATCCGGTTGTGGTATATTTGGAATTTCTACTGTTGTAGATTAGCATAACCCCTTGGGGCCTCTAAACGGG | Construction of pUDR490 |
| **14278** | longT7pGG_gADE2-3 | GCCGGGAATTTAATACGACTCACTATAGGAATTTCTACTGTTGTAGATCCGGTTGTGGTATATTTGGAATTTCTACTGTTGTAGATTAGCATAACCCCTTGGGGCCTCTAAACGGG | Construction of pUDR491 |
| **14279** | longT7pGGG_gADE2-3 | GCCGGGAATTTAATACGACTCACTATAGGGAATTTCTACTGTTGTAGATCCGGTTGTGGTATATTTGGAATTTCTACTGTTGTAGATTAGCATAACCCCTTGGGGCCTCTAAACGGG | Construction of pUDR492 |

| | | | |
|---|---|---|---|
| **11751** | pUDC191_Fr2_fw | GGGTCATCACGGCTCATCATGC | Backbone amplification for *in vivo* plasmid construction |
| **12378** | Backbone pGGKd017 REV | ATTGCGACGAATTGCCACG | |
| **14584** | pUD-gRNA_fw | GCGATCACAGACATTAACCCACA | gRNA cassette amplification for *in vivo* plasmid construction or expression from linear template |
| **14585** | pUD-gRNA_rv | GCAGTCATCCGAGCGTGTATTG | |
| **6815** | KanMX (new)_Fw | ATGTTGGACGAGTCGGAATC | Amplification of backbone for split plasmid selection |
| **9340** | E.coli- rv | GCACGGTTATCCACAGAATCAG | |
| **2398** | RV pMA-RQ | GTTTCGCCACCTCTGACTTG | |
| **12097** | YTK_kanR rv | TTAGAAAAACTCATCGAGCATC | |
| **Construction of T7RNAP mutant strains (IMX2030, IMX2031, IMX2032) and overexpression strain (IME475)** | | | |
| **14284** | sgRNA_T7RNP | TGCGCATGTTTCGGCGTTCGAAACTTCTCCGCAGTGAAAGATAAATGATCTGCCAGCCAGCGCACCTGCAGTTTTAGAGCTAGAAATAGCAAGTTAAAATAAG | Construction of plasmid pUDR506 |
| **6005** | p426 CRISP rv | GATCATTTATCTTTCACTGCGGAGAAG | |
| **6006** | p426 CRISP fw | GTTTTAGAGCTAGAAATAGCAAGTTAAAATAAGGCTAGTC | |
| **14379** | T7rep-P266L_fw | TCGAACTCGCACCTGAATACGCTGAGGCTATCGCAACACGTGCAGGTGCGCTGGCTGGCATCTCTCTGATGTTCCAACCTTGCGTAGTTCCTCCTAGGCCGTGGACTGGCATTACTGGTG | Repair oligos for IMX2030 construction |
| **14380** | T7rep-P266L_rv | CACCAGTAATGCCAGTCCACGGCCTAGGAGGAACTACGCAAGGTTGGAACATCAGAGAGATGCCAGCCAGCGCACCTGCACGTGTTGCGATAGCCTCAGCGTATTCAGGTGCGAGTTCGA | |
| **14665** | T7rep-R276K_fw | TCGAACTCGCACCTGAATACGCTGAGGCTATCGCAACACGTGCAGGTGCGCTGGCTGGCATCTCTCCGATGTTCCAACCTTGCGTAGTTCCTCCTAAGCCGTGGACTGGCATTACTGGTG | Repair oligos for IMX2031 construction |
| **14666** | T7rep-R276K_rv | CACCAGTAATGCCAGTCCACGGCTTAGGAGGAACTACGCAAGGTTGGAACATCGGAGAGATGCCAGCCAGCGCACCTGCACGTGTTGCGATAGCCTCAGCGTATTCAGGTGCGAGTTCGA | |
| **14667** | T7rep-P266L.R276K_fw | TCGAACTCGCACCTGAATACGCTGAGGCTATCGCAACACGTGCAGGTGCGCTGGCTGGCATCTCTCTGATGTTCCAACCTTGCGTAGTTCCTCCTAAGCCGTGGACTGGCATTACTGGTG | Repair oligos for IMX2032 construction |
| **14668** | T7rep-P266L.R276K_rv | CACCAGTAATGCCAGTCCACGGCTTAGGAGGAACTACGCAAGGTTGGAACATCAGAGAGATGCCAGCCAGCGCACCTGCACGTGTTGCGATAGCCTCAGCGTATTCAGGTGCGAGTTCGA | |
| **13710** | T7in_rv | GAGTAACACCGTAAGCCAGC | Diagnostic T7RNAP SNPs |
| **14738** | T7RNP.266P_fw | GCTGGCTGGCATCTCTCC | |
| **14739** | T7RNP.266L_fw | GCTGGCTGGCATCTCTCT | |
| **14740** | T7RNP.276R_fw | ACCTTGCGTAGTTCCTCCTAG | |
| **14741** | T7RNP.276K_fw | ACCTTGCGTAGTTCCTCCTAA | |
| **10753** | TDH3 sc prom fw Ytoolkit | AAGCATCGTCTCATCGGTCTCAAACGCGAATATATACTAGCGTTGAATGTTAG | Construction of pUDE911 |
| **10768** | TEF1 sc term rv Ytoolkit | TTATGCCGTCTCAGGTCTCACAGCGGTATCACCATAGATTTCGAAAC | |

| | | | |
|---|---|---|---|
| **Construction of plasmids for *HIS4*, *PDR12* and *CAN1* editing via Cas12a nuclease** | | | |
| **17328** | gHIS4-4sGGG | AAACTTCTCCGCAGTGAAAGATAAATGATCGGGAATTTCTACTGTTGTAGATGCATCTTGGCTAGCAATGAAATTTCTACTGTTGTAGATTTTTTTTGTTTTTTATGTCTGCTGCCAATG | Construction of pUDR715 |
| **17329** | gPDR12-3sGGG | AAACTTCTCCGCAGTGAAAGATAAATGATCGGGAATTTCTACTGTTGTAGATGCACAAAGAATCAATATGGAATTTCTACTGTTGTAGATTTTTTTTGTTTTTTATGTCTGCTGCCAATG | Construction of pUDR716 |
| **17330** | gCAN1-4sGGG | AAACTTCTCCGCAGTGAAAGATAAATGATCGGGAATTTCTACTGTTGTAGATCCACATATCTTCAACGCTGAATTTCTACTGTTGTAGATTTTTTTTGTTTTTTATGTCTGCTGCCAATG | Construction of pUDR717 |
| **17331** | gCAN1-3sGGG | AAACTTCTCCGCAGTGAAAGATAAATGATCGGGAATTTCTACTGTTGTAGATAAGGTACTGAACTAGTTGGAATTTCTACTGTTGTAGATTTTTTTTGTTTTTTATGTCTGCTGCCAATG | Construction of pUDR718 |
| **2398** | RV pMA-RQ | GTTTCGCCACCTCTGACTTG | Diagnostic and Sanger sequencing |
| **10325** | ConR1_rv | AGTCATCCGAGCGTGTATTG | |
| **gDNA oligos for T7RNAP-mediated gRNA expression** | | | |
| **15089** | T7p-GGG-gADE2-3-DR-T7t_fw | GCCGGGAATTTAATACGACTCACTATAGGGAATTTCTACTGTTGTAGATCCGGTTGTGGTATATTTGGAATTTCTACTGTTGTAGATTAGCATAACCCCTTGGGGCCTCTAAACGGG | Cas12a ADE2-3 gDNA (Long T7p, DR, short T7t) |
| **15090** | T7p-GGG-gADE2-3-DR-T7t_rv | CCCGTTTAGAGGCCCCAAGGGGTTATGCTAATCTACAACAGTAGAAATTCCAAATATACCACAACCGGATCTACAACAGTAGAAATTCCCTATAGTGAGTCGTATTAAATTCCCGGC | |
| **15091** | T7p-GGG-gADE2-3-T7t_fw | GCCGGGAATTTAATACGACTCACTATAGGGAATTTCTACTGTTGTAGATCCGGTTGTGGTATATTTGGTAGCATAACCCCTTGGGGCCTCTAAACGGG | Cas12a ADE2-3 gDNA (Long T7p, short T7t) |
| **15092** | T7p-GGG-gADE2-3-T7t_rv | CCCGTTTAGAGGCCCCAAGGGGTTATGCTACCAAATATACCACAACCGGATCTACAACAGTAGAAATTCCCTATAGTGAGTCGTATTAAATTCCCGGC | |
| **15093** | T7p-GGG-gADE2-3-DR_fw | GCCGGGAATTTAATACGACTCACTATAGGGAATTTCTACTGTTGTAGATCCGGTTGTGGTATATTTGGAATTTCTACTGTTGTAGAT | Cas12a ADE2-3 gDNA (Long T7p, DR) |
| **15094** | T7p-GGG-gADE2-3-DR_rv | ATCTACAACAGTAGAAATTCCAAATATACCACAACCGGATCTACAACAGTAGAAATTCCCTATAGTGAGTCGTATTAAATTCCCGGC | |
| **15252** | lT7p-GGG-gADE2-3_fw | GCCGGGAATTTAATACGACTCACTATAGGGAATTTCTACTGTTGTAGATCCGGTTGTGGTATATTTGG | Cas12a ADE2-3 gDNA (Long T7p) |
| **15253** | lT7p-GGG-gADE2-3_rv | CCAAATATACCACAACCGGATCTACAACAGTAGAAATTCCCTATAGTGAGTCGTATTAAATTCCCGGC | |
| **15254** | T7p-GGG-gADE2-3-DR_fw | TAATACGACTCACTATAGGGAATTTCTACTGTTGTAGATCCGGTTGTGGTATATTTGGAATTTCTACTGTTGTAGAT | Cas12a ADE2-3 gDNA (Short T7p, DR) |
| **15255** | T7p-GGG-gADE2-3-DR_rv | ATCTACAACAGTAGAAATTCCAAATATACCACAACCGGATCTACAACAGTAGAAATTCCCTATAGTGAGTCGTATTA | |
| **15290** | lT7p-GGG-gADE2-3_fw | GCCGGGAATTTAATACGACTCACTATAGGGAATTTCTACTGTTGTAGATCCGGTTGTGGTATATTTGG | Cas9 ADE2.y gDNA (Short T7p) |
| **15291** | lT7p-GGG-gADE2-3_rv | CCAAATATACCACAACCGGATCTACAACAGTAGAAATTCCCTATAGTGAGTCGTATTAAATTCCCGGC | |

| | | | |
|---|---|---|---|
| **16477** | lT7p-GGG-gADE2-3-T7t_fw | GCCGGGAATTTAATACGACTCACTATAGGGAATTTCTACTGTTGTAGATCCGGTTGTGGTATATTTGGTAGCATAACCCCTTGGGGCCTCTAAACGGGTCTTGAGGGGTTTTTTG | Cas12a ADE2-3 gDNA (Long T7p, long T7t) |
| **16478** | lT7p-GGG-gADE2-3-T7t_rv | CAAAAAACCCCTCAAGACCCGTTTAGAGGCCCCAAGGGGTTATGCTACCAAATATACCACAACCGGATCTACAACAGTAGAAATTCCCTATAGTGAGTCGTATTAAATTCCCGGC | |
| **16745** | lT7p-GGG-ADE2.y_fw | GCCGGGAATTTAATACGACTCACTATAGGGACTTGAAGATTCTTTAGTGTGTTTTAGAGCTAGAAATAGCAAGT | PCR-derived Cas9 ADE2.y gDNA (Long T7p) |
| **16746** | struct-gRNA_rv | GCACCACCGACTCGGTGCCACTTTTTCAAGTTGATAACGGACTAGCCTTATTTTAACTTGCTATTTCTAGCTCTAAAAC | |
| **16479** | lT7p-GGG-gCAN1-4-DR_fw | GCCGGGAATTTAATACGACTCACTATAGGGAATTTCTACTGTTGTAGATCCACATATCTTCAACGCTGAATTTCTACTGTTGTAGAT | Cas12a CAN1-4 gDNA (Long T7p, DR) |
| **16480** | lT7p-GGG-gCAN1-4-DR_rv | ATCTACAACAGTAGAAATTCAGCGTTGAAGATATGTGGATCTACAACAGTAGAAATTCCCTATAGTGAGTCGTATTAAATTCCCGGC | |
| **16481** | lT7p-GGG-gHIS4-4-DR_fw | GCCGGGAATTTAATACGACTCACTATAGGGAATTTCTACTGTTGTAGATGCATCTTGGCTAGCAATGAAATTTCTACTGTTGTAGAT | Cas12a HIS4-4 gDNA (Long T7p, DR) |
| **16482** | lT7p-GGG-gHIS4-4-DR_rv | ATCTACAACAGTAGAAATTTCATTGCTAGCCAAGATGCATCTACAACAGTAGAAATTCCCTATAGTGAGTCGTATTAAATTCCCGGC | |
| **16483** | lT7p-GGG-gPDR12-3-DR_fw | GCCGGGAATTTAATACGACTCACTATAGGGAATTTCTACTGTTGTAGATGCACAAAGAATCAATATGGAATTTCTACTGTTGTAGAT | Cas12a PDR12-3 gDNA (Long T7p, DR) |
| **16484** | lT7p-GGG-gPDR12-3-DR_rv | ATCTACAACAGTAGAAATTCCATATTGATTCTTTGTGCATCTACAACAGTAGAAATTCCCTATAGTGAGTCGTATTAAATTCCCGGC | |
| **16699** | lT7p-GGG-gCAN1-3-DR_fw | GCCGGGAATTTAATACGACTCACTATAGGGAATTTCTACTGTTGTAGATAAGGTACTGAACTAGTTGGAATTTCTACTGTTGTAGAT | Cas12a CAN1-3 gDNA (Long T7p, DR) |
| **16700** | lT7p-GGG-gCAN1-3-DR_rv | ATCTACAACAGTAGAAATTCCAACTAGTTCAGTACCTTATCTACAACAGTAGAAATTCCCTATAGTGAGTCGTATTAAATTCCCGGC | |
| **Repair oligos for gene deletions** | | | |
| **10155** | Ade2repairFw | ATGGATTCTAGAACAGTTGGTATATTAGGAGGGGGACAATTGGGACGTATGATTGTTGAGCTTGTCAAAGCACAAAAGTTAGAAACTGTCGGTTACGAAGCTTATCTAGAAAACAAGTAA | Deletion of *ADE2* |
| **10156** | Ade2repairRv | TTACTTGTTTTCTAGATAAGCTTCGTAACCGACAGTTTCTAACTTTTGTGCTTTGACAAGCTCAACAATCATACGTCCCAATTGTCCCCCTCCTAATATACCAACTGTTCTAGAATCCAT | |
| **10226** | 10226_His4 repair Fw | CGATGTGTGTTGTACATACATAAAAATATCATAGCACAACTGCGCTGTGTAATAGTAATACGAACACTAACGAAAATAATATGTATATATACATATATATATCAAACAAAATACAGTCTT | Deletion of *HIS4* |
| **10227** | 10227_His4 repair Rv | AAGACTGTATTTTGTTTGATATATATATGTATATATACATATTATTTTCGTTAGTGTTCGTATTACTATTACACAGCGCAGTTGTGCTATGATATTTTTATGTATGTACAACACACATCG | |
| **7042** | PDR12 Repair Oligo FW | AAAATTGAAAATAAAAATTGTGTGTTAAACCACGAAATACAAATATATTTGCTTGCTTGTTTTTTTATTAATAAGAACAATAACAATAAATCTGTAAACCTTTTTTTTAAGTGAAAATTA | Deletion of *PDR12* |

| | | | |
|---|---|---|---|
| **7043** | PDR12 Repair Oligo RV | TAATTTTCACTTAAAAAAAAGGTTTACAGATTTATTGTTATTGTTCTTATTAATAAAAAA ACAAGCAAGCAAATATATTTGTATTTCGTGGTTTAACACACAATTTTTATTTTCAATTTT | |
| **5614** | Oligo CAN 1 fw | TATGAGGGTGAGAATGCGAAATGGCGTGGAAATGTGATCAAAGGTAATAAAACGTCA TATTGCTATGCCTTTTTTTTTTTTTGTTTTTACAGGAGTTAAGAAGTCTGAAGAACTCTG AAA | Deletion of *CAN1* |
| **5615** | Oligo CAN 1 rv | TTTCAGAGTTCTTCAGACTTCTTAACTCCTGTAAAAACAAAAAAAAAAAAAGGCATAGC AATATGACGTTTTATTACCTTTGATCACATTTCCACGCCATTTCGCATTCTCACCCTCATA | |
| **<u>Diagnostic PCR of gene deletions</u>** | | | |
| **3603** | ade2-AMDS fw3 | TCTAAGTACATCCTACTATAACAATC | Diagnostic PCR for *ADE2* deletion |
| **2681** | ADE2dsHRV | GGACACTTATATGTCGAGCAAGA | |
| **10228** | His4 del check Fw | AACAGCCGTGGAATCGTTGC | Diagnostic PCR for *HIS4* deletion |
| **10229** | His4 del check Rv | TGGGTAGTGCCTTGTGATCC | |
| **3997** | PDR12 KO fw | CTTTCGGCTATGGCTCATAC | Diagnostic PCR for *PDR12* deletion |
| **254** | PDR12 - CTRL RV | GGAAGATAATAGGCGAGCTC | |
| **2496** | FW-conf-upstrm | CGGGAGCAAGATTGTTGTG | Diagnostic PCR for *CAN1* deletion |
| **5821** | CAN1 PAGE fw | CGAATCAGGGAATCCCTTTTTG | |

#### Table S2 - Cloning free genome editing techniques

Comparison of cloning-free methods for genome editing in *Saccharomyces cerevisiae*

| Name | N of targets: Efficiency | Cas nuclease[a] | gRNA expression (promoter; cassette construction) | Reference |
|---|---|---|---|---|
| **pMEL** | 1: 25-75%<br>3: 0% | *Sp*Cas9 (genome integrated) | *SNR52p*;<br>double-stranded annealed oligos, *in vivo* plasmid assembly | (2) |
| **CAM** | 1: 27-100%<br>3: 64% | *Sp*Cas9 (genome integrated) | *SNR52p*;<br>PCR generated from preassembled plasmid, *In vivo* plasmid assembly | (3) |
| **pRCC** | 1: ~60%<br>2: ~15% | *Sp*Cas9 (from multicopy plasmid) | *SNR52p*;<br>PCR generated from template cassette, *in vivo* plasmid assembly | (4) |
| **scCRISPR/Cpf1** | 1: 80%<br>3: 32% | *Fn*Cas12a (from centromeric plasmid) | *SNR52p*;<br>PCR generated from oligo template, *In vivo* plasmid assembly | (5) |
| **Lightning GTR-CRISPR** | 4: 96%<br>6: 60% | *Sp*Cas9 (from multicopy plasmid) | *SNR52p*;<br>PCR generated from template cassette, golden gate plasmid assembly | (6) |
| **gRNA-TES** | 1: 67-100% | *Sp*Cas9 (from centromeric plasmid) | *SNR52p*;<br>PCR generated from template cassette | (7) |
| **gEL DNA** | 1: 96%<br>2: 63%<br>4: 3% | *Fn*Cas12a (from multicopy plasmid) | *T7p*;<br>double-stranded annealed oligos | This study |

[a] Cas protein is constitutively expressed.

#### Table S3 – Sequence and secondary structure of the gDNAs used in this study

Features of gRNAs for Cas12a used in this study and prediction of lowest free-energy RNA folding. The predicted sequence of the gDNA folding consists of three Gs, 19-bp direct repeat (DR:AAUUUCUACUGUUGUAGAU) and 19-bp spacer sequence (underlined). Query sequence for the crRNA secondary structure prediction consists of 19-bp direct repeat (DR) and 25-bp spacer sequence as in Swiat *et al.,* (8). Self-folding free energies are indicated at the right of the structure prediction. The colour-code indicates base-pair probabilities according to the legend below the table.

| **gRNA target** | **crRNA sequence (shorter <u>19-bp</u> spacer underlined)** | **gRNA for T7 polymerase (3G-DR-19bp spacer)** | **gRNA used in Swiat *et al.* (DR-25bp spacer)** |
|---|---|---|---|
| ADE2-3 | <u>CCGGUUGUGGUAUAUUUGG</u>UGUGGA | 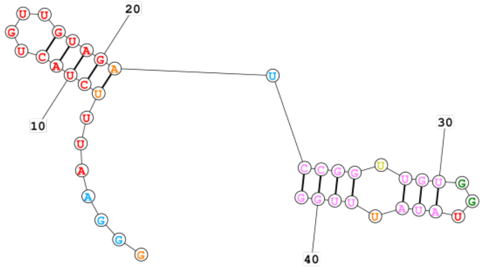 -9.6 | 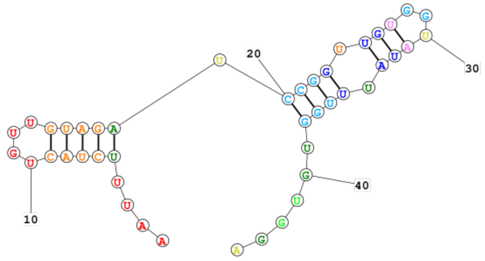 -9.6 |
| HIS4-4 | <u>GCAUCUUGGCUAGCAAUGA</u>ACAGAG | 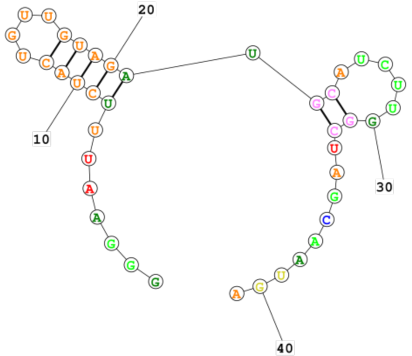 -8.6 | 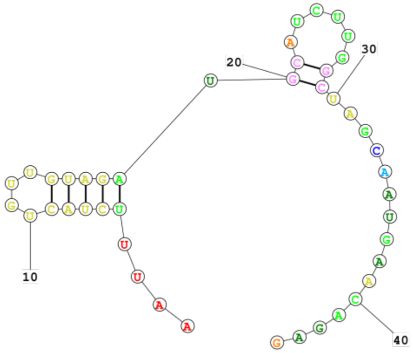 -8.6 |

| | | | |
|---|---|---|---|
| PDR12-3 | [GGG]-DR-GCACAAAGAAUCAAUAUGGGUGUCA | 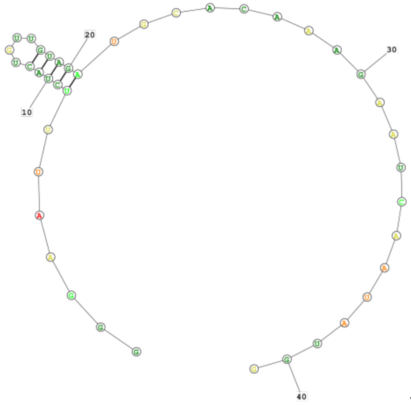 -5.7 | 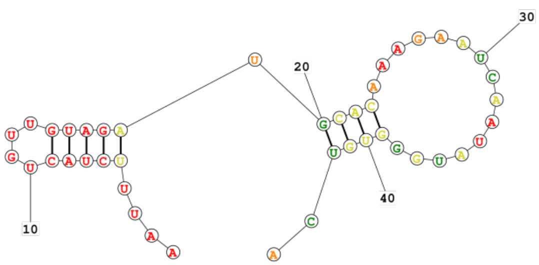 -9.4 |
| CAN1-4 | [GGG]-DR-CCACAUAUCUUCAACGCUGUUAUCU | 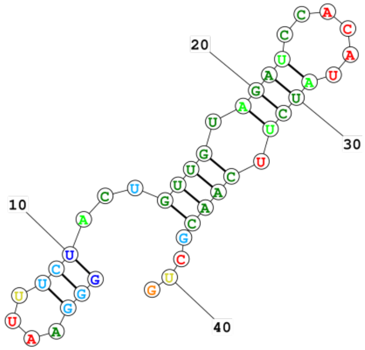 -9.1 | 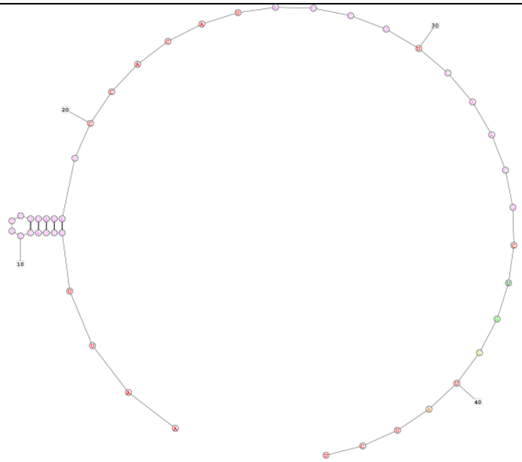 -5.7 |

| | | | |
|---|---|---|---|
| CAN1-3 | [GGG]-DR-AAGGUACUGAACUAGUUGGUAUCAC | 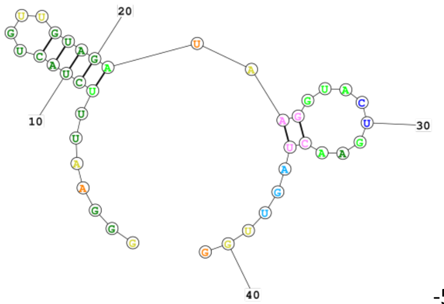 -5.9 | 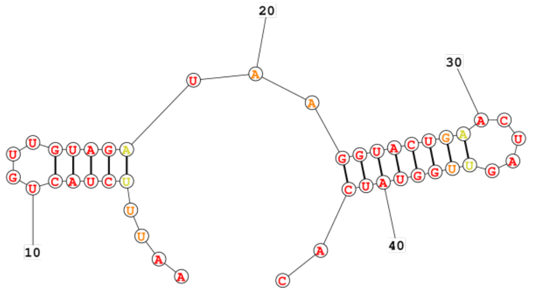 -13.1 |

Legend:

Probability >= 99%
99% > Probability >= 95%
95% > Probability >= 90%
90% > Probability >= 80%
80% > Probability >= 70%
70% > Probability >= 60%
60% > Probability >= 50%
50% > Probability
